# High-throughput binding affinity measurements for mutations spanning a transcription factor-DNA interface reveal affinity and specificity determinants

**DOI:** 10.1101/2020.06.22.165571

**Authors:** Arjun K. Aditham, Craig J. Markin, Daniel A. Mokhtari, Nicole V. DelRosso, Polly M. Fordyce

## Abstract

Transcription factors (TFs) bind regulatory DNA to control gene expression, and mutations to either TFs or DNA can alter binding affinities to rewire regulatory networks and drive phenotypic variation. While studies have profiled energetic effects of DNA mutations extensively, we lack similar information for TF variants. Here, we present STAMMP (Simultaneous Transcription Factor Affinity Measurements via Microfluidic Protein Arrays), a high-throughput microfluidic platform enabling quantitative characterization of hundreds of TF variants simultaneously. Measured affinities for ∼210 mutants of a model yeast TF (Pho4) interacting with 9 oligonucleotides (>1,800 *K*_d_s) reveal that many combinations of mutations to poorly conserved TF residues and nucleotides flanking the core binding site alter but preserve physiological binding, providing a mechanism for mutations in *cis* and *trans* to rewire networks without insurmountable evolutionary penalties. Moreover, biochemical double-mutant cycles across the TF-DNA interface reveal molecular mechanisms driving recognition, linking sequence to function.

## Introduction

Regulation of gene expression is critical for proper organismal development and response to environmental stimuli. This regulation is accomplished primarily by transcription factor (TF) proteins that bind specific regulatory DNA sequences in the genome to activate or repress gene expression (Mitsis, et al., 2020). Reflecting this central role in cellular function, mutations to either DNA regulatory sequences or TF proteins have profound effects across evolution and medicine. Mutations that alter TF-DNA interactions in *cis* (DNA binding sites) or in *trans* (diffusible factors, *e.g*. TFs) drive organismal evolution by rewiring transcriptional networks and generating phenotypic variation (Signor and Nuzhdin, 2018; Wong, et al., 2018). In medicine, the majority of disease-associated mutations in humans are found within regulatory DNA (Maurano et al., 2012; Nishizaki, et al., 2020), and mutations in TFs often lead to cancers (Lambert M, et al., 2018; Lee and Young, 2013) and developmental disorders (Lee and Young, 2013; Barrera, et al., 2016). A quantitative and predictive understanding of how mutations affect TF/DNA binding interactions would therefore have broad impacts for understanding biology.

The probability of TF occupancy at a given genomic site is typically modeled as a function of the Gibbs free energy of binding (ΔG) for that TF/DNA sequence combination and the effective concentration of free TFs in the nucleus (Foat, et al., 2006, Kim and O’Shea, 2008, Segal, et al., 2008; Gertz et al., 2009; Weirauch, et al., 2015; Zhao and Stormo, 2015; Le, et al., 2018). As a result, differences in binding site affinity and available TF concentrations can modulate the strength and timing of transcriptional programs (Crocker, et al., 2016). While much effort has focused on determining the highest affinity sites for a given TF (Chen, et al., 2016; Jolma, et al., 2010; Zykovich, et al., 2009), non-specific and low-affinity sites in the genome often play essential functional roles and are evolutionarily conserved (Farley, et al., 2015; Crocker, et al., 2016; Kribelbauer, et al., 2019). Given that even subtle changes in affinity can alter transcription (Meinhardt, et al., 2013; Rajkumar, et al., 2013; Le, et al., 2018), understanding how TF and DNA mutations affect binding requires the ability to measure affinities with high accuracy over a wide energetic range.

Many prior efforts have comprehensively characterized how variation in underlying DNA sequence alters TF binding energies for up to 10^6^ DNA variants in parallel (Le, et al., 2018). These studies have established that typical TF-DNA binding energies span nearly 2-3 orders of magnitude (∼3 kcal/mol) (Nutiu, et al., 2011; Kribelbauer, et al., 2019), that mutations to nucleotides within or flanking the “core” consensus site can alter affinities over a wide range (Le, et al., 2018; Maerkl and Quake, 2007), and that measured binding energies can predict levels of induction and rates of transcription *in vivo* (Gaudet and Mango, 2002; Aow, et al., 2013).

By contrast, far less is known about how mutations throughout a TF protein alter DNA binding, even though binding energies depend on the sequences of both the underlying DNA *and* the TF. This lack of data likely stems from the technical challenges associated with high-throughput expression, purification, and quantitative biophysical characterization of large TF mutant libraries. As a result, previous studies have typically focused on limited sets of residues, such as those that directly contact the DNA sequence (Ferreiro, et al., 2008; Maerkl and Quake, 2009; Wang, et al., 2017), regulate dimerization of protein partners (Voronova and Baltimore, 1990), or are associated with human disease (Barrera, et al., 2016). While these studies have confirmed that mutations at strongly conserved residues typically dramatically alter DNA binding affinity, the effects of mutations at poorly conserved residues remain understudied (Meinhardt, et al., 2013). Large quantitative datasets describing the effects of mutations throughout and surrounding TF binding domains on DNA binding could reveal mechanistic insights not easily inferred from static crystal structures and provide critical information about binding interfaces for TFs recalcitrant to crystallization efforts (Fuxreiter, et al., 2011). In addition, such datasets would also provide an invaluable resource for optimizing computational methods designed to predict mutational effects for precision medicine (Ng and Henikoff, 2003; Adzhubei, et al., 2010; Choi, et al., 2012) and create synthetic transcriptional circuits.

Here, we report the development of STAMMP (Simultaneous Transcription Factor Affinity Measurements via Microfluidic Protein Arrays), a microfluidic platform capable of measuring binding affinities for >1500 TF mutants simultaneously interactive with a DNA sequence of interest in a single experiment. As a first application, we systematically quantify mutational effects on DNA binding affinity and specificity for Pho4, a homodimeric basic helix-loop-helix (bHLH) TF in yeast that binds a 5’-CACGTG-3’ E-box motif to drive gene expression in response to phosphate starvation (Ogawa, et al., 2000). We measured more than 1,800 binding affinities (*K*_d_s or ΔGs) for ∼210 Pho4 mutants interacting with oligonucleotides containing mutations within and flanking the core E-box site and found that a large fraction (>70%) of residue mutations have statistically significant effects on DNA binding. Strikingly, of the more than 1800 pairs of TF and DNA mutations assayed, nearly 70% of mutations that altered but did not ablate binding occurred at nucleotides outside of the core binding site and at residues that do not contact DNA nucleotides. These combinations of mutations in *cis* and in *trans* could tune affinities to enable network rewiring without large-scale deleterious effects, suggesting that previously unexplored residues and mutations may play an unexpected role in evolution of new function. Finally, biochemical double mutant cycles across the TF-DNA interface provide mechanistic insight into the interactions that govern recognition, quantifying the energetic contributions of contacts predicted from the crystal structure and revealing additional residues required for specificity. We anticipate that STAMMP and the presented data will be broadly useful for future development of quantitative models designed to link TF sequence to structure and function.

## Results

### STAMMP enables high-throughput characterization of the functional effects of TF mutations on DNA binding

High-throughput functional characterization of large numbers of TF mutants requires the ability to recombinantly express and purify hundreds of TFs in parallel. To accomplish this task, STAMMP affinity measurements take place within a microfluidic device containing 1,568 valved reaction chambers, each of which contains a “plasmid” compartment and a “binding reaction” compartment **(Figs. 1A, S1)** (Fordyce, et al., 2010; Maerkl and Quake, 2007). Three sets of valves control fluid flow within and between reaction chambers: “neck” valves **(Fig. 1A, green)** separate plasmid and binding compartments, “sandwich” valves **(Fig. 1A, red)** physically sequester reaction chambers from one another to prevent cross-contamination, and “button” valves **(Fig. 1A, blue)** in the binding compartment enable selective surface patterning within the device and trap macromolecular binding interactions at equilibrium for quantitative affinity measurements.

**Figure 1.**
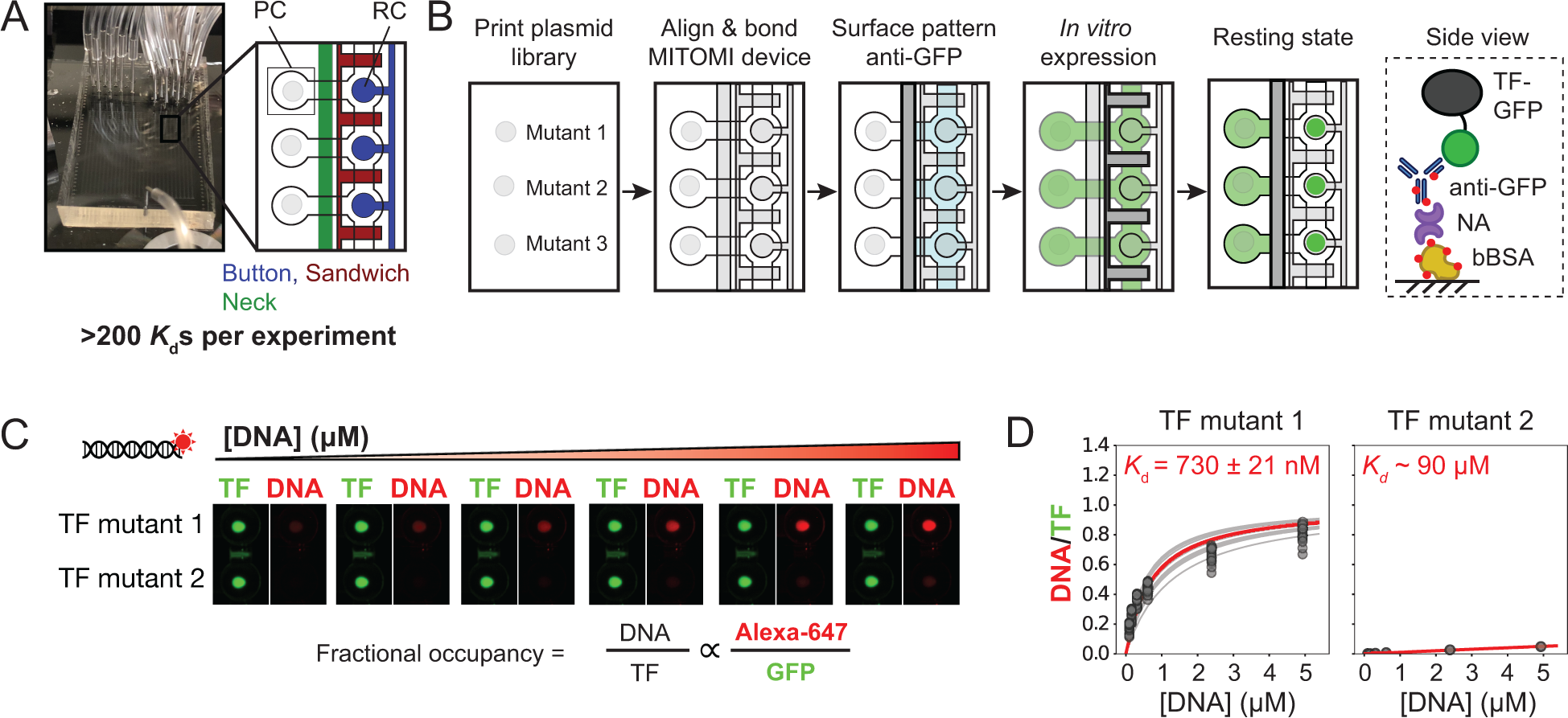
STAMMP: microfluidic platform for high-throughput TF expression and assays. **(A)** Photograph of MITOMI device showing plasmid (labeled as “PC”) and binding reaction compartments (labeled as “RC”) and annotated pneumatic valves (“button”, “sandwich”, and “neck”). **(B)** Workflow for device surface functionalization and *in situ* expression and purification of TF libraries and final schematic of components used for surface-immobilization of eGFP-tagged TFs (right). **(C)** Workflow illustrating iterative introduction, equilibration, and trapping of TF-bound fluorescently-labeled DNA to quantify concentration-dependent binding; final ratios of DNA and TF fluorescence are proportional to fractional occupancy. Image contrast was enhanced to improve visibility for display purposes. **(D)** Langmuir isotherms fit to DNA/TF fluorescence intensity ratios as a function of free DNA concentration yield interaction binding affinities. Example raw binding data (grey markers and fit lines) and calculated median fit curves (red lines) for 2 mutants are shown at right; *K*_d_ values represent median values ± SEM for statistical replicates.

To program each reaction chamber with a specific TF variant, microfluidic devices are aligned to spotted plasmid arrays **(Fig. 1B)**. Each plasmid encodes expression of a TF variant fused to a C-terminal monomeric eGFP tag (Zacaharias, et al., 2002), with the identity of each mutant encoded by its position in the array. After alignment, device surfaces are specifically patterned with anti-eGFP antibodies (anti-GFP) beneath the “button” valve and coated elsewhere with biotinylated bovine serum albumin (bBSA) to prevent non-specific adsorption. All TF variants are then expressed in parallel via introduction of cell-free extract into each chamber and incubation of the device for 1 hour at 37°C. After expression, TFs are purified via recruitment to antibody-patterned surfaces **(Figs. 1B, S2)**. Subsequent closing of the “button” valves allows extensive washing and trypsin digestion to remove any nonspecifically bound TFs and remove cell-free expression components while protecting surface-immobilized protein; the final eGFP fluorescence intensity in each chamber reports on the amount of immobilized TF **(Fig. 1B,C)**.

To measure full binding isotherms for all expressed TFs interacting with a particular oligonucleotide sequence, we iteratively introduce fluorescently-tagged DNA duplexes at multiple concentrations across all chambers **(Figs. 1C, S2)**. After allowing reactions to come to equilibrium, we again close the “button” valves in each chamber, thereby trapping all bound TF-DNA complexes. After washing, we image all reaction chambers across the device to quantify TF and DNA intensities and estimate fractional occupancies **(Fig. 1C)**. We then fit estimated fractional occupancies as a function of the effective concentration of free DNA in solution (calculated from intensities using per-chamber calibration curves, **Fig. S3**) to a Langmuir isotherm to extract binding affinities (*K*_d_s) and relative differences in binding affinities (ΔΔGs) for all TFs in parallel (Fig. 1D).

### A library of Pho4 mutants designed to probe mechanisms of binding site recognition

Basic helix-loop-helix (bHLH) TFs are highly abundant in eukaryotes, representing the third most abundant TF structural class in the human genome (Lambert, et al., 2018). Pho4 is one of eight bHLHs in *S. cerevisiae* (Chen and Lopes, 2007), and prior work has established that Pho4 binds the E-box core motif (5’-CACGTG-3’) element to drive gene expression in response to phosphate starvation. In addition to this well-characterized and inducible biological function, Pho4 has a well-characterized DNA binding interface (Shimizu, et al., 1997; Cave, et al., 2000) and a wealth of prior biophysical data describing how mutations to the underlying DNA sequence affect affinity (Maerkl and Quake, 2007; Fordyce, et al., 2010; Le, et al., 2018). Comparisons between the Pho4 crystal structure and those of other bHLH proteins (Shimizu, et al., 1997; Brownlie, et al., 1997; Parraga, et al., 1998) also show structural similarity, suggesting that Pho4 can serve as a model for understanding how bHLH specificity is encoded in protein sequence.

The DNA binding domains (DBDs) of all bHLH TF superfamily members contain a helical basic region (which directly contacts DNA) and two additional helices separated by an unstructured loop of variable length **(Fig. 2A)**. Four residues within the basic region directly contact nucleobases within the major groove of the E-box recognition site and are strongly conserved throughout organisms, including yeast and humans (R252, H255, E259, and R263 in Pho4); additional residues within the loop contact the DNA backbone **(Fig. 2A, Fig. S4)**. Pho4 binds DNA as a homodimer **(Fig. 2B)**, with conserved hydrophobic residues in the helix 1 and helix 2 regions promoting stable dimerization **(Fig. 2A)**.

**Figure 2.**
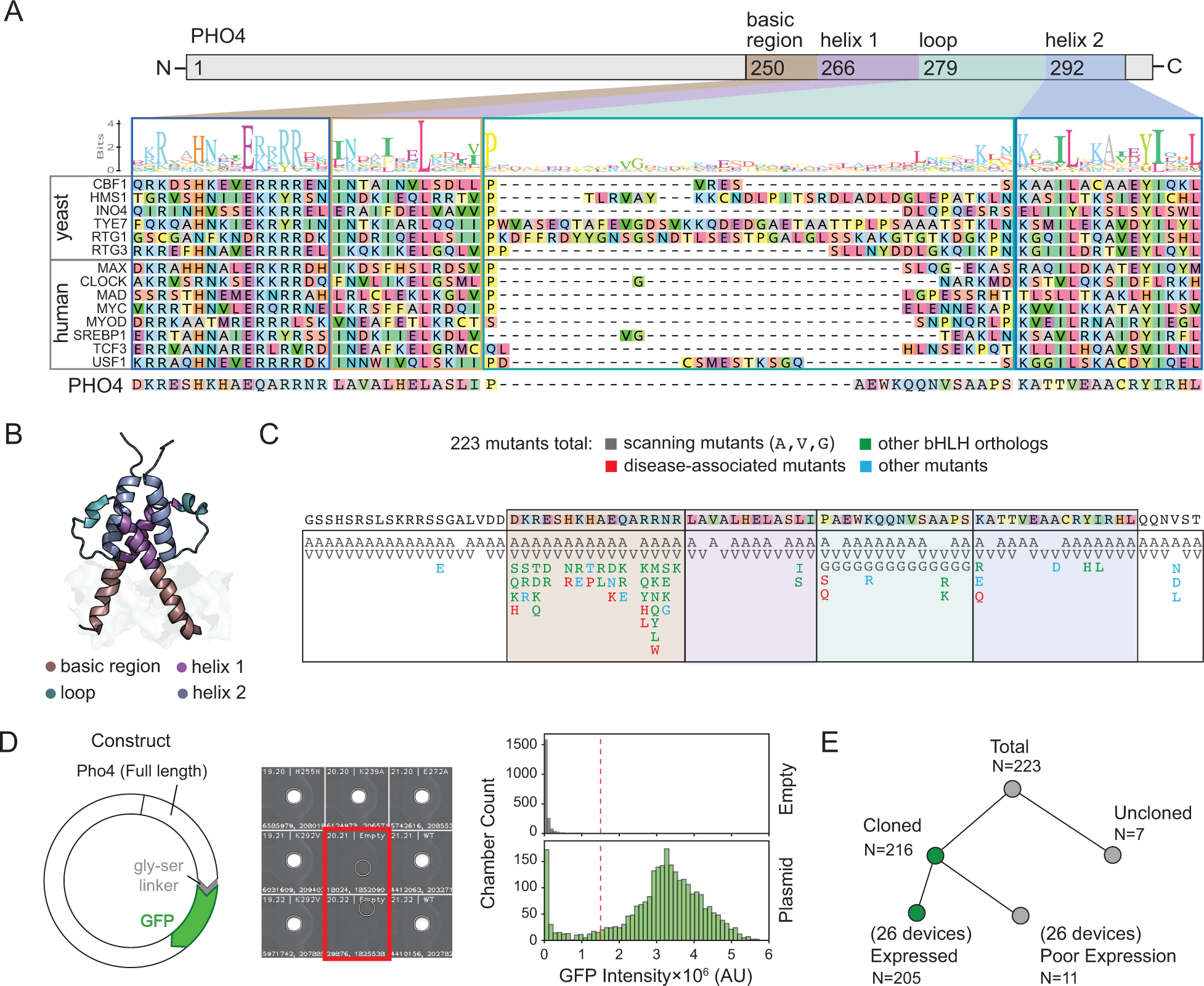
Pho4 mutant library design and expression. **(A)** Protein sequence alignment for bHLH proteins across yeast and humans. **(B)** Pho4 structure (PDB: 1A0A) with regions within DNA-binding domain labeled. **(C)** Pho4 library mutants categorized by scan, presence in other bHLH proteins, representation in disease variants, and other types of mutations (charge modulation, *etc*.). **(D)** Plasmid map of full-length Pho4 construct fused to c-terminal GFP tag (left); sample image of 9 of 1568 device chambers showing measured fluorescence intensities for chambers containing Pho4-eGFP plasmid and empty chambers (red) (middle); and measured intensities for all 1568 chambers in a representative device (right). **(E)** Binary tree summarizing mutant outcomes for all 223 mutants in the library.

To systematically probe how residues throughout and immediately surrounding the DNA binding domain (DBD) contribute to binding affinity and specificity, we generated a library of 223 Pho4 mutants comprised of systematic substitutions, variants from other bHLH orthologs, known disease-associated variants from human orthologs, and a selection of mutants designed to probe mechanisms of binding **(Fig. 2C)**. For systematic substitutions, we introduced alanine and valine residues at each site, thereby ablating side chains and substituting a hydrophobic moiety, respectively. To probe disease-associated and evolutionary variants, we introduced mutations from the human bHLH MAX previously observed in tumor samples and made substitutions to residues observed in orthologs across ascomycetes, humans, and *C. elegans* (DeMasi, et al., NAR, 2011) at the corresponding Pho4 positions. Finally, we tested the effects of altering the electrostatic charge of amino acids on Pho4 binding by substituting charged residues for those with the opposite charge or by substituting uncharged residues for positively charged residues.

After on-chip expression, immobilization, and purification, expressed TFs were visible as high-intensity spots within each reaction chamber in the eGFP channel, while chambers lacking plasmid typically showed no fluorescence **(Fig. 2D)**. A small fraction of chambers lacking plasmid yielded observable intensities, providing a sensitive readout of potential cross-contamination between chambers that was used to set a lower threshold for identifying chambers with successful expression. Overall, 216 of 223 mutants (∼97%) successfully expressed at least once **(Fig. 2E)** and 205 of 223 mutants (∼92%) expressed consistently across the 26 experiments in this study **(Fig. 2E)**.

### Most mutations throughout Pho4 have statistically significant effects on DNA binding

First, we tested which residues contribute to Pho4 DNA binding by quantifying the effects of amino acid mutations on measured binding affinity for 5’-C CACGTG A-3’, a cognate DNA sequence with medium-affinity flanking nucleotides chosen to enhance the ability to observe beneficial and deleterious effects on binding (Le, et al., 2018). We generated fluorescently-labeled DNA constructs by annealing an Alexa647-labeled primer to a universal sequence at the 3’ end of each DNA and extending constructs using Klenow polymerase fragment **(Fig. 3A)**. Measured binding affinities were highly reproducible between experimental replicates **(Fig. S5)**, allowing reliable determination of mutations that both increased and decreased affinity relative to wildtype Pho4. To estimate affinities for deleterious variants with *K*_d_ values above concentrations probed experimentally, we globally fit all binding curves to a Langmuir isotherm with a single shared saturation value and individually-fit *K*_d_ values (Fordyce, et al., 2010; Fordyce, et al., 2012; Nguyen, et al., 2019). Extracted *K*_d_ and ΔΔG values for all mutants across experiments were highly reproducible over an energetic range of ∼4 kcal/mol (*K*_d_: *r*^2^ = 0.96, RMSLE = 0.22 nM; ΔΔG: *r*^2^ = 0.96, RMSE = 0.31 kcal/mol) **(Figs. 3B,C, S5)**. We designated mutants with measured ΔΔG values not statistically significantly different (using a two-tailed t-test with a Bonferroni correction) from those for a mutant lacking DNA-binding activity (A299D, which likely disrupts the dimerization interface) as binding-deficient (see Methods, **Fig. S6, S7**).

**Figure 3.**
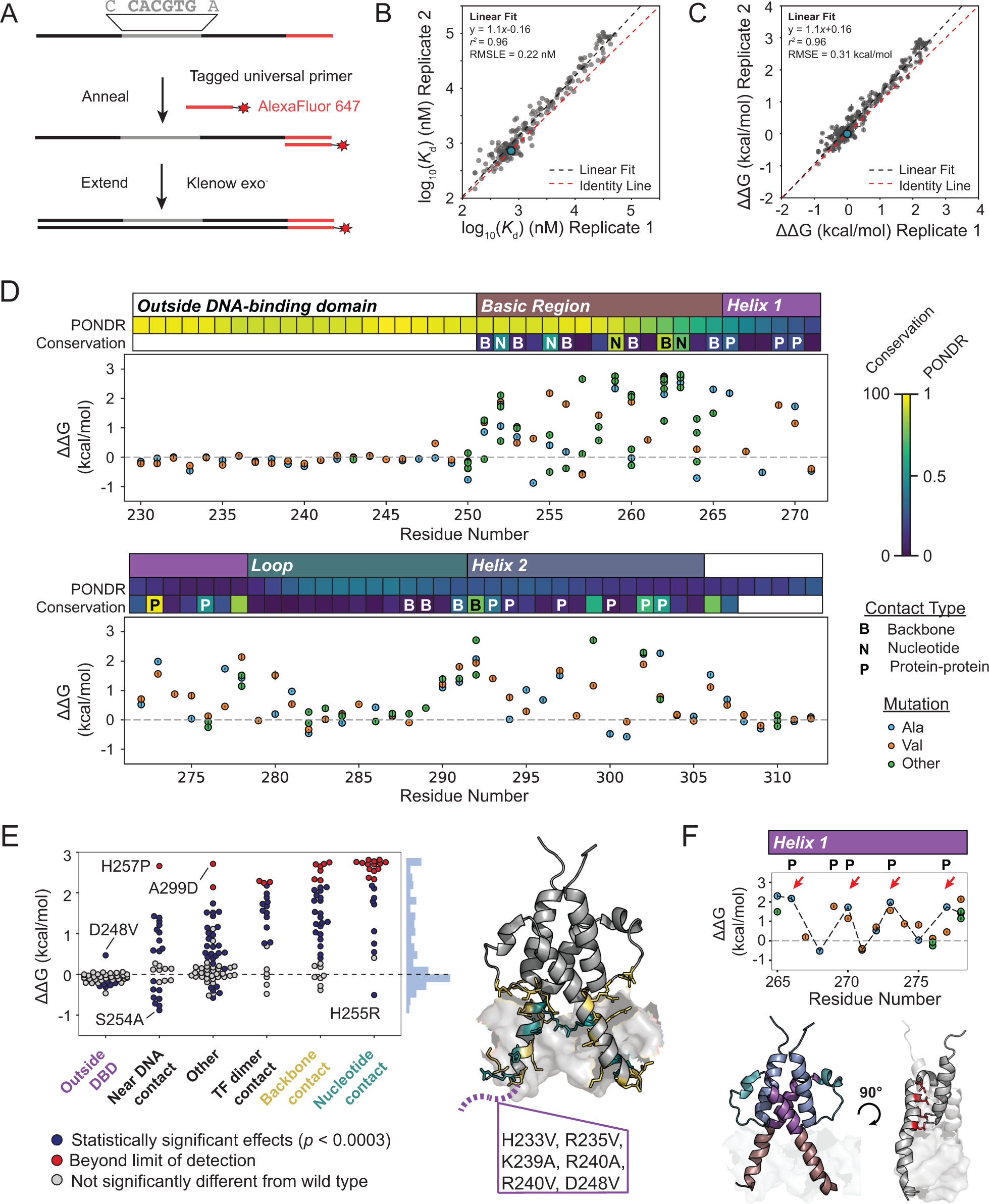
Many mutations throughout the Pho4 DBD alter DNA binding. **(A)** Workflow for creating fluorescently-labeled DNA duplexes for binding assays. **(B)** Pairwise comparison of measured *K*_d_ values (median ± SEM) for all TF mutants interacting with a 5’-CCACGTGA-3’ sequence assayed across two separate devices (n=190); WT Pho4 is shown in blue. Grey and red dashed lines indicate a linear regression and the identity line, respectively. Affinity measurements were log-transformed and fitted to a linear model; RMSLE was then calculated from log-transformed values. **(C)** Pairwise comparison of measured DDGs (median ± SEM) for Pho4 mutants; WT Pho4 is shown in blue. Grey and red dashed lines indicate a linear regression and the identity line, respectively. **(D)** Median DDG of binding (± SEM) for all Pho4 mutants interacting with a 5’-CCACGTGA-3’ sequence; known crystal structure contacts, amino acid conservation, and predicted intrinsic disorder are shown. **(E)** Median DDG of binding for all mutations binned by interaction with DNA based on crystal structure (right) and labeled by statistical difference from wildtype Pho4 and background binding. **(F)** Periodicity of effects on DNA binding (top) and indicated positions on Pho4 crystal structure (bottom). Data reflects median DDG (± SEM) for DNA binding; dashed line passes through alanine mutants.

A large fraction of mutations to residues within and outside of the DNA binding domain had statistically significant effects on DNA binding (∼56%), including at positions not in contact with DNA **(Fig. 3D)**. As expected, mutating residues outside of the DNA binding domain generally had little to no effect, while mutations to nucleotide- and backbone-contacting residues typically reduced binding to below the limit of detection **(Figs. 3D,E)**. Despite poor sequence conservation of the bHLH loop region **(Fig. 2A)**, mutations to residues in the loop close to either helix 1 or helix 2 had strong effects on binding, likely due to either proximity to the DNA molecule and known backbone contacting residues or via disruption of essential secondary structure **(Fig. S8)**. Although mutations to strongly conserved residues tended to be more deleterious overall (*r*^*2*^ = 0.41), effects were strongly mutation dependent, with different substitutions at a given position yielding changes in binding affinities that varied by ∼2 kcal/mol **(Figs. S8, S9)**. PROVEAN, a computational algorithm designed to predict the functional effects of mutations, was moderately successful at predicting whether mutations with large energetic effects were damaging or benign using a discrete cut-off (AUC = 0.82); however, the overall correlation between the magnitudes of predicted and observed effects was relatively poor (*r*^*2*^ = 0.35) **(Figs. S9)**. Taken together, these results indicate that phylogeny alone is not sufficient to predict observed mutational effects on affinity.

Multiple mutations to residues near DNA contacts and outside of the DNA-binding domain significantly *increased* DNA affinity, including H255R, which corresponds to a hotspot cancer variant in the human MAX protein (Cerami, et al., 2012; Gao, et al., 2013; Wang, et al., 2017). These results suggest that this particular variant may exact its deleterious effects by increasing nonspecific binding to DNA in the genome. This is also consistent with prior suggestions that TFs are evolutionarily selected to have moderate affinities *in vivo* to enable dynamic transcriptional responses to changing nuclear TF concentrations (Gaudet and Mango, 2002; Wang, et al., 2017; Kribelbauer, et al., 2019) **(Figs. 3D,E)**. The removal of positively charged residues in the N-terminal intrinsically disordered tail **(Fig. S10)** yielded small but statistically significant increases in DNA binding affinity, suggesting that charged residues outside the DBD might play a role in DNA binding in Pho4 as reported for other TFs (Vuzman and Levy, 2010; Vuzman and Levy, 2012; Shammas, 2017).

### Systematic mutagenesis coupled to high-resolution affinity measurements can reveal secondary structure

Many TFs, including bHLHs, include large disordered regions that are thought to assume a folded 3D structure only upon binding DNA and therefore pose technical challenges for crystallization. We therefore sought to assess whether systematic mutational scans coupled with quantitative affinity measurements could reveal clues regarding secondary structures in the absence of crystal structures or NMR data. Observed periodic effects of mutations within helix 1 mapped to the reference crystal structure (Shimizu, et al., 1997) revealed that deleterious mutations lie at the protein dimerization interface while benign mutations can be found at the solvent-exposed exterior **(Fig. 3F)**. Moreover, valine substitutions tended to be more deleterious compared to alanine mutations at the DNA interface (likely due to their steric bulk) while alanine substitutions were disfavored compared to valine mutations at the protein-protein interface (likely due to disruptions in hydrophobic packing) **(Fig. S11)**. Scanning mutagenesis with amino acid substitutions of varied biochemical properties coupled to high-resolution energetic measurements may therefore have utility for guiding computational structural predictions and for explaining mechanisms of residue function within a host protein.

### Comparing effects of TF mutations across oligonucleotide sequences reveals residues involved in modulating affinity and specificity

TF mutations may alter transcriptional regulation by uniformly modulating DNA binding affinity (*i.e*. altering affinity to all DNA sequences equally), modulating DNA specificity (*i.e*. differentially affecting affinity for some DNA sequences), or a combination of the two. To distinguish between affinity and specificity effects, we systematically measured effects of TF mutations on binding to 8 additional DNA sequences comprised of single-residue mutations within the core 5’-CACGTG-3’ E-box motif as well as the first flanking nucleotide on either side **(Fig. 4A, Figs. S12-14)**. Consistent with the known Pho4 position weight matrix (PWM) and a variety of previous reports, we observed that mutations to the core E-box motif significantly reduced affinity and a 5’ C flanking nucleotide was preferred (Fisher and Goding, 1992; Maerkl and Quake, 2007; Le, et al., 2018; Spivak and Stormo, 2012). Measured affinities were ∼6-fold weaker than seen previously (Maerkl and Quake, 2007), likely due to low levels of nonspecific adsorption of expressed TFs on chamber walls (see Methods); ΔΔGs are a relative measurement and are therefore unaffected.

**Figure 4.**
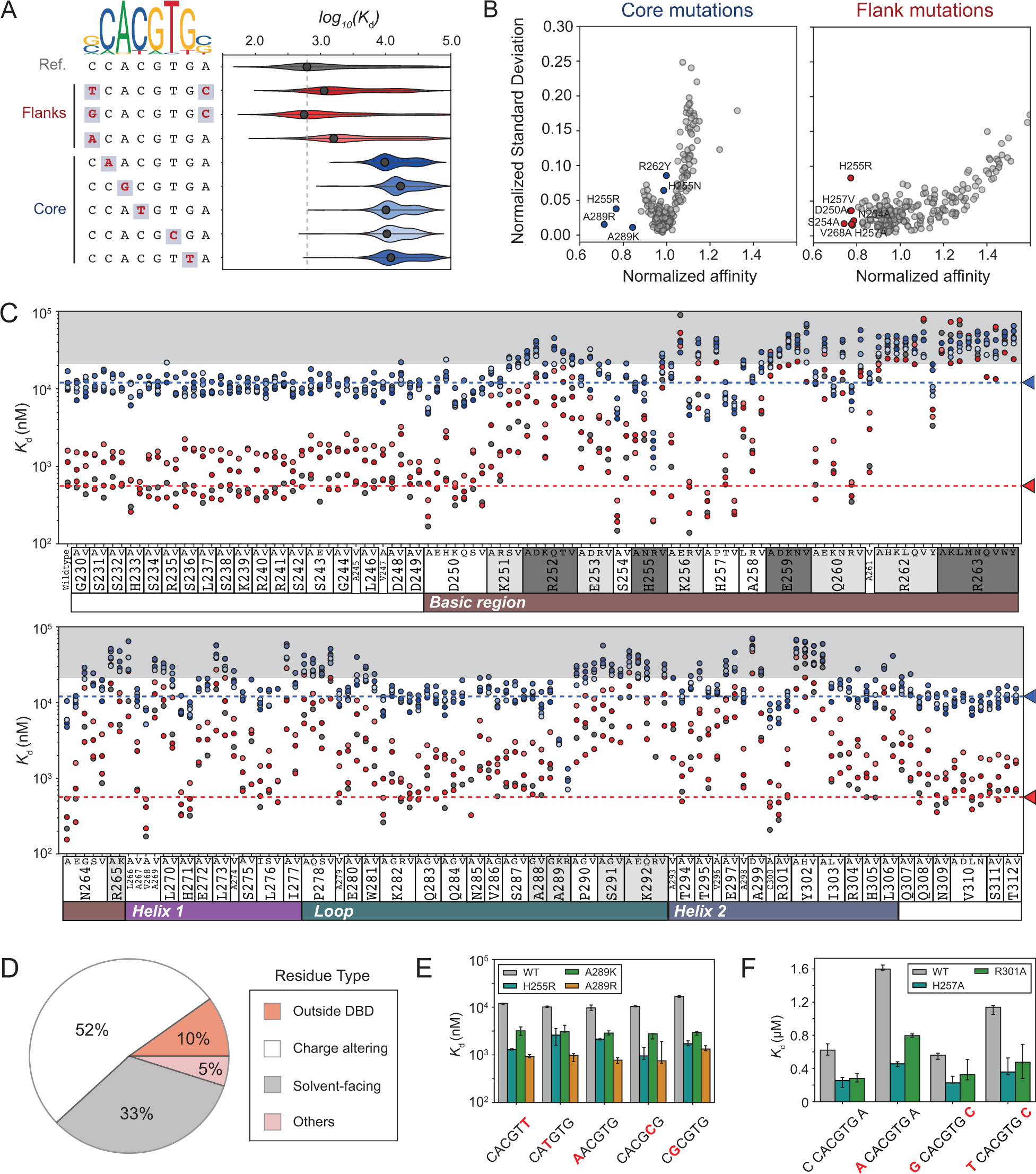
Effects of Pho4 mutations on DNA affinity and specificity. **(A)** PWM logo and list of DNA sequences studied alongside effects on binding affinity for all Pho4 variants studied. Black datapoints reflect affinity of wildtype Pho4 for DNA sequence; vertical dashed line indicates median affinity of WT Pho4 for the reference oligo 5’-C CACGTG A - 3’. **(B)** Normalized standard deviation versus normalized affinities for all Pho4 mutants indicating mutants with effects on affinity and/or specificity. **(C)** Measured median binding affinities across experimental replicates for all TF mutations interacting with all oligonucleotides. Blue arrow indicates median wildtype Pho4 affinity for the low-affinity 5’-C CACGTT A-3’ sequence; red arrow indicates median wildtype Pho4 affinity for the DNA sequence (5’-G CACGTG C-3’). Points are colored by oligo identify, as in Fig. 4A. **(D)** Proportions of mutations that statistically significantly enhance DNA binding affinity. **(E)** Binding affinities for WT Pho4 and H255R, A289K, and A289R mutants (median affinity ± SEM). (F) Binding affinities for Pho4 H257A (Helix 1) and R301A (Helix 2) interacting with DNA sequences containing flanking nucleotide mutations (median affinity ± SEM).

To identify TF mutants with affinity or specificity effects across multiple DNA sequences, we quantified the spread in measured DNA binding affinities (standard deviation) normalized by the median affinity for each oligo (log_10_(*K*_d_)_i,j_/<log10(*K*_d_)_j_> where *i* is a particular TF mutation and *j* is a particular oligonucleotide mutation) **(Fig. 4B)**. Overall, standard deviations increase as measurements diverge towards very high or low affinity (particularly for core mutations), reflecting increased measurement noise towards the dynamic range limits of the assay. Several mutants (H255R, A289R, and A289K) increase the median affinity without increasing the standard deviation, likely by positioning positively charged residues in proximity to the negatively charged DNA backbone **(Figs. 4B, S15)**. Several mutations to critical DNA-contacting residues, R262Y and H255N, both of which are observed in the phylogenetic record (Kim, et al., 1995; Shimizu, et al., 1997; del Olmo Toledo, et al., 2018; Figure 2A) appeared to increase the variance in measured affinities, suggesting that these mutations may primarily alter specificity **(Fig. 4B)**.

Measured affinities plotted as a function of mutated residue position for all 1853 unique TF-DNA mutant pairs reveal several striking trends **(Fig. 4C)**. *In vivo*, WT Pho4 binds both the reference E-box sequence (5’-CACGTG-3’) and a known biologically-relevant low-affinity site (5’-CACGTT-3’) (Barbaric, et al., 1996; Zhou and O’Shea, 2011); we therefore used these values as guidelines for establishing a range of physiologically relevant binding **(Fig. 4C)**. For WT Pho4 and for mutations outside of the DBD, changes to less favorable flanking nucleotides or mutations in the core binding site typically reduced affinities by ∼4- and 20-fold, respectively, preserving low-to-medium affinity binding. For many residues within the DBD (and particularly for those that make direct contacts with DNA), the combined effects of TF and DNA mutations reduced binding to that of the dynamic range floor **(Fig. 4C)**. However, a large number of DBD mutants retained the ability to bind oligonucleotides containing flanking sequence mutations with a range of physiologically-relevant binding affinities. Overall, 128 of the 151 mutants with altered binding relative to the WT protein retained the ability to bind at least some oligonucleotides at levels above background. Moreover, some of these mutants have differential effects for different flanking sequences. This ability to ‘tune’ binding affinity via combinations of TF and DNA mutations provides a potential mechanism for rewiring regulatory networks *in trans* while avoiding unwanted pleiotropic effects.

### Differences in electrostatics and helical propensity modulate binding affinity

Closer examination of residues that significantly increased DNA binding affinity revealed that many appeared to introduce an additional positive charge near the DNA (∼52%, as for H255R, A289K, and A289R) **(Figs. 4D,E, S16)**. However, a large fraction did so without altering charge at DNA-contacting residues, including H257A and R301A **(Figs. 4D,F)**. Approximately 33% of mutations found to enhance affinity were primarily alanine or valine substitutions at solvent-facing residues **(Figs. 4D, 5A, S16)**, highlighting the role of non-contacting and distal amino acid residues in dictating binding affinity and spurring mechanistic questions regarding how they exert their effects. Prior studies have revealed that helix 1 of Pho4 and other bHLH TFs are unstructured in the absence of DNA and adopt a primarily helical conformation upon binding cognate DNA (Cave, et al., 2000; Sauvé, et al., 2004). Prior work has shown that modulating the helical propensity of TF sequences can tune binding affinity for DNA sequences containing the E-box motif by stabilizing the helical conformation of the TF (Kunne, et al., 1997; Turner, et al., 2004) **(Fig. 5B)**. To test if changes in helical propensity were sufficient to explain observed differences in measured binding affinity for alanine and valine mutants, we compared measured changes in Gibbs free energy of binding with previously determined changes in Gibbs free energy for helix formation (O’Neil and DeGrado, 1990) for all residues near DNA-contacting residues **(Fig. 5A)** and found that these were highly correlated (*r*^*2*^ = 0.58; RMSE = 0.70 kcal/mol) **(Fig. 5C)**. As expected, measured ΔΔG values for mutations at nucleotide-contacting residues in helical regions were dominated by changes in hydrogen bonding and only weakly correlated with predicted changes in helical propensity (*r*^*2*^ = 0.27; RMSE = 2.0 kcal/mol; **Fig S17A)**, and mutations to residues in the loop region, which remains unstructured regardless of DNA binding, showed no correlation (*r*^*2*^ = 0.0089; RMSE = 1.2 kcal/mol; **Fig S17B)**. Together, these results strongly suggest that mutations in helical regions that modulate the equilibrium between unstructured and helical conformations can tune transcription factor binding.

**Figure 5.**
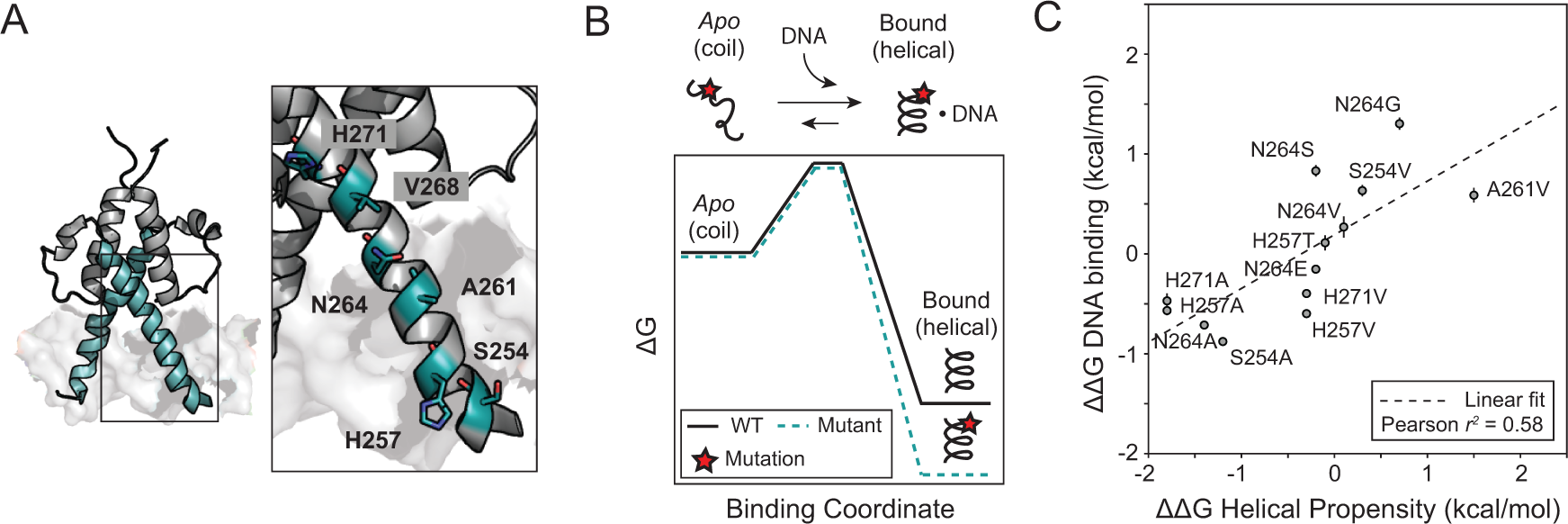
Solvent-facing residues modulate affinity by tuning helical propensity. **(A)** Crystal structure of Pho4 with basic region and Helix 1 highlighted (cyan); right, inset of solvent-facing residues in this region. **(B)** Schematic reaction coordinate diagram for WT (solid black line) and mutated Pho4 (cyan dashed line) in which the mutant construct has a higher helical stability. **(C)** Measured change in DNA binding affinity (ΔΔG, median ± SEM) for the 5’-C CACGTG A-3’ reference oligonucleotide *vs*. previously measured changes in helical propensity for individual residue substitutions (O’Neil and DeGrado, 1990); dashed line indicates linear regression.

### Double mutant cycles across the TF-DNA interface can reveal intermolecular interactions required for recognition

Beyond simply cataloging effects, this set of 1,853 quantitative affinities provides a unique opportunity to measure the strength of inter-molecular interactions across the TF-DNA interface via biochemical double mutant cycles (Horovitz, 1996; Horovitz, et al., 2019). In general, we expect that mutating residue(s) essential for recognizing a particular nucleotide will be relatively *less* deleterious upon that nucleotide’s mutation. As a simplified example, consider a TF that binds a DNA sequence such that residue 1 does not directly contact any nucleotides but residue 2 makes a direct contact with nucleotide 2 **(Fig. 6A)**. For residue 1, mutating this residue will have the same relative effect on affinity regardless of the oligonucleotide sequence. For residue 2, mutating this residue will significantly reduce binding to the WT oligonucleotide but have a less deleterious effect for an oligonucleotide in which nucleotide 2 has been mutated. Systematic comparisons of the relative effects on affinity for all TF mutants for each DNA sequence can therefore provide a comprehensive method for revealing functional interactions required for specific recognition and binding.

**Figure 6:**
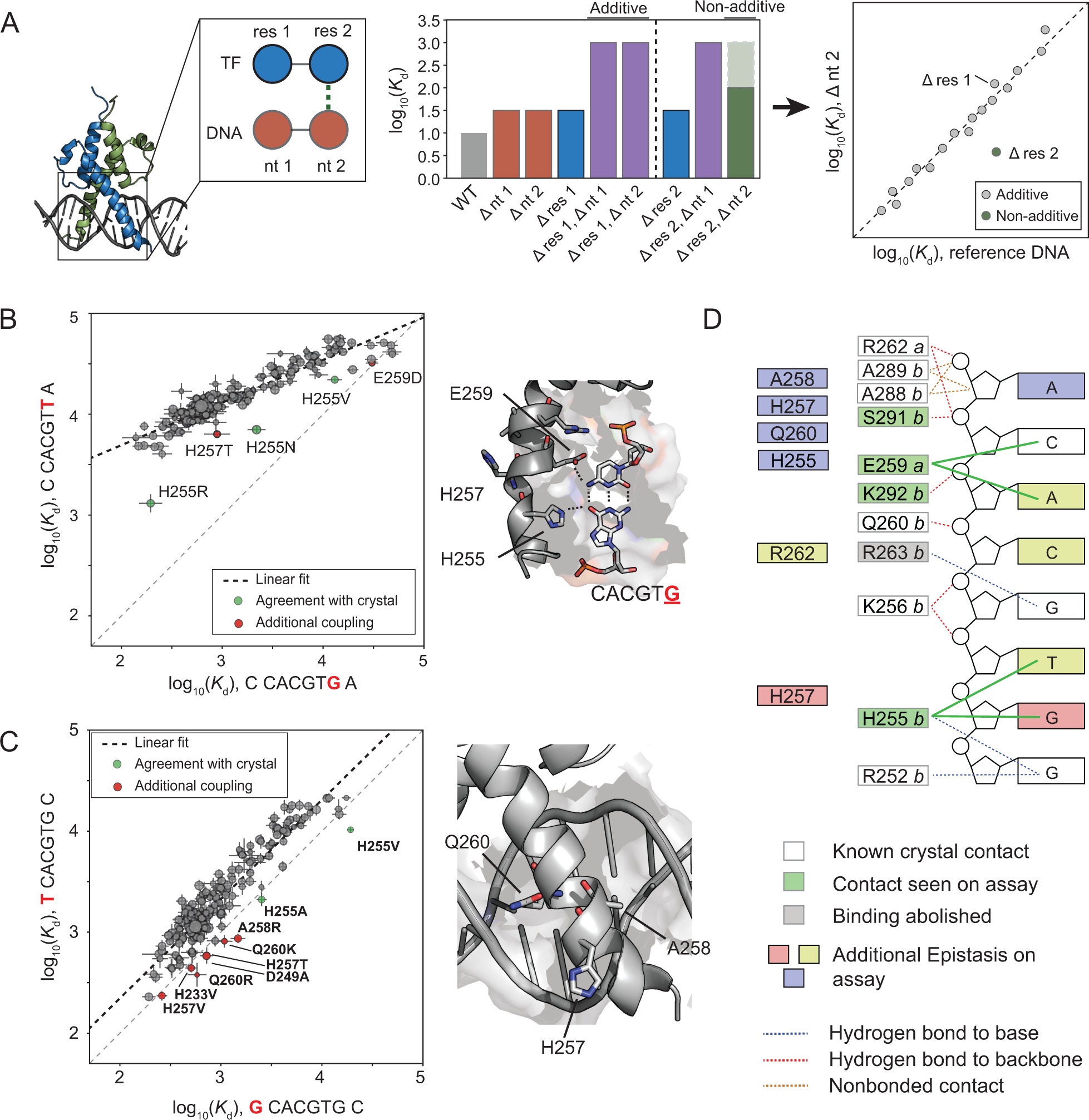
Pairwise affinity comparisons reveal determinants of binding specificity. **(A)** Example cartoon illustrating how epistasis between Pho4 and DNA mutations can be detected by comparing energetic effects of TF mutations across different DNA sequences. **(B)** Pairwise comparison between measured binding affinities (log_10_(*K*_d_), median ± SEM) for Pho4 mutants interacting with a low-affinity mutant sequence (5’-CCACGT<u>T</u>A-3’) *vs*. the medium affinity reference sequence (5’-CCACGTGA-3’) (left); inset of crystal structure with outlier TF residues labeled. **(C)** Pairwise comparison between measured binding affinities (log_10_(*K*_d_), median ± SEM) for 2 DNA sequences with 5’ flanking nucleotide mutations (left); inset of crystal structure indicating residues H257, A258, and Q260 (right). **(D)** Comparison between identified energetic couplings identified in this study and known crystallographic contacts.

As a first application of this approach, we compared affinities measured for the Pho4 mutant library interacting with the biologically relevant low-affinity 5’-CCACGTTA-3’ site and the WT reference E-box sequence (5’-CCACGTGA-3’). The crystal structure for Pho4 bound to a DNA duplex containing a 5’-CCACGTGT-3’ suggests a direct contact between the H255 residue and the final G nucleotide within the E-box **(Fig. 6B, inset)**, predicting that mutating H255 should have differential effects for sequences containing a G or a T at this position. As expected, measured affinities were weaker overall for the mutant DNA sequence, with most mutants having similar effects (**Figs. 6B, S18-19**; *r*^*2*^ = 0.84). However, several mutants were significantly less deleterious for binding the mutated DNA sequence than the cognate DNA sequence, including the expected H255 mutants (H255R, H255N, H255V), H257T, and E259D. Inspection of measured concentration-dependent binding curves confirmed energetic non-additivity for combined TF and DNA mutations **(Fig. S19D)**. These additional observed energetic effects can be explained by previously observed structural contacts: the E259 residue contacts the C nucleotide base-paired with the G on the opposite strand, and H257 is a solvent-facing residue on the same helix as H255, suggesting that H257T might alter the conformation of DNA-contacting residues H255 and E259 or the overall helicity of the TF to modulate binding. These results therefore establish that double mutant cycles across the TF-DNA interface via STAMMP represent a high-throughput approach for detecting relevant functional interactions required for recognition.

### Double mutant cycles reveal TF residues required to specify flanking nucleotide preferences

For Pho4, preferences for nucleotides flanking the core consensus site play a critical role in dictating *in vivo* occupancies, rates of gene activation, and levels of induction (Aow, et al., 2013; Gordân, et al., 2013; Rajkumar, et al., 2013; Le, et al., 2018). However, the residues responsible for mediating this specificity remain unknown. While the crystal structure of Pho4 suggested that DNA contacting residues R252 and H255 mediate specificity (Shimizu, et al., 1997), mutagenesis studies of Pho4 studies have implicated residues outside of the DNA binding domain (Fisher and Goding, 1992) and demonstrated that mutations in orthologous TFs corresponding to the Pho4 A258 residue can alter flanking nucleotide specificity (Beltran et al., 2004).

To investigate the origins of flanking nucleotide specificity, we compared measured affinities for a sequence in which the first 5’ flanking nucleotide was mutated to a T (5’-T CACGTG C-3’) with those for a 5’-G CACGTG C-3’ variant **(Figs. 6C, S20)**. Although measured affinities for the mutant oligonucleotide were slightly weaker for nearly all mutants, a small set of mutants bound the flanking 5’ T nucleotide more tightly. This set included multiple mutations to H255, consistent with previous findings that this residue helps mediate flanking nucleotide specificity (Shimizu, et al., 1997). However, we also detected enhanced binding for the A258R, H257T, Q260R, and Q260K variants (Shimizu, et al., 1997; Beltran, et al., 2004) **(Figs. 6C, S20D)**, establishing that mutations within the DNA binding domain can alter specificity **(Fig. 6C)**. These substitutions include both aliphatic and positively charged residues, suggesting that introducing steric bulk at the DNA interface (as for A to R and Q to K substitutions) may modulate flanking nucleotide specificity (Fisher and Goding, 1992).

Repeating this analysis for mutations throughout the core binding site recapitulated many contacts predicted by the known crystal structure **(Fig 6D, S19-24)**. For mutations that abolish binding across all oligonucleotides (*e.g*. mutations at R263 and most back-bone contacting residues) **(Figs. 6D, S21-23)**, double mutant cycle analysis cannot quantify epistasis; however, these critical contacts are typically easily identified from phylogeny. In many other cases, double mutant cycle analysis reveals functional consequences that could not be predicted purely from static structures. Here, R262Y, located at a known DNA backbone contact, was the only mutant at either position 262 or 263 still capable of binding DNA, consistent with phylogenetic data showing that the human bHLH family member SREBP1 contains a tyrosine at the orthologous position and that this mutation enhances conformational flexibility to allow binding to varied E-box sequences (del Olmo Toledo, et al., 2018; Parraga, et al., 1998; **Fig. S25**). In addition, this analysis identified specificity-altering mutations (e.g. H255N and R262Y) that cannot be predicted from a crystal structure crystallized in the presence of a single DNA sequence. High-throughput thermodynamic measurements can therefore complement and supplement traditional structural analysis for identifying macromolecular contacts.

## Discussion

Here, we present STAMMP, which enables quantitative measurements of DNA binding affinities across >1,500 TF variants within a single experiment. Using this system, we measured absolute and relative binding energies for ∼210 Pho4 mutants interacting with multiple DNA sequences, including substitutions at positions spanning the core binding site and the 5’ and 3’ flanking nucleotides. In total, 76% (n=163) of TF mutations led to statistically significant differences in DNA binding; of these, 80% (n=133) preserved binding above background levels. A large fraction of these residues does not directly contact DNA (either nucleobases or the backbone), suggesting that many poorly conserved positions that were previously unexplored experimentally may play critical roles in specifying and remodeling transcriptional responses.

Mutations in *cis* have long been considered the primary drivers of evolutionary variation (Nelson, et al., 2013; Wong, et al., 2018), as TF mutations can have pleiotropic effects on cellular function that limit evolvability and diversification of genetic networks (Signor and Nuzhdin, 2018.). These conclusions have been further bolstered by experimental evidence that mutations to DNA-contacting residues or core consensus nucleotides often ablate binding (Maerkl and Quake, 2009; De Masi, et al., 2011), creating large fitness penalties that pose evolutionary barriers. Here, the ability to assess many mutations in parallel and measure even subtle differences in affinity suggests that effects of mutations in *trans* could be compensated for by concomitant changes in nucleotides flanking *cis-*regulatory elements to preserve transcriptional responses (Beltran, et al., 2004; Rogers and Bulyk, 2018). In addition, the observation that mutations may stabilize helical conformations to promote DNA binding suggests that these residues could allow formation of permissive binding intermediates (Bloom, et al., 2010; Gong, et al., 2013; Jalal, et al., 2019; McKeown, et al., 2014; Starr, et al., 2017). The higher affinity binding of these permissive binding intermediates could allow TFs to acquire and accommodate additional TF or DNA mutations that would otherwise reduce binding to non-physiological levels, making otherwise inaccessible evolutionary trajectories feasible.

Beyond providing evolutionary insights, we demonstrate here that STAMMP measurements can provide information that can be used to better predict structures for TFs refractory to crystallization. Obtaining structures for TFs is often technically challenging because many TFs assume a folded conformation only upon interacting with a consensus binding site and contain large, intrinsically disordered domains. Reflecting this, of 1639 predicted TFs in the human genome, only 185 (∼11%) have crystal structures available (Lambert, et al., 2018). Of these structures, most are solely composed of the DBD alone and are crystallized only in the presence of a single high-affinity DNA sequence. STAMMP can provide critical functional information about residues within intrinsically disordered regions previously shown to be important for DNA binding and dynamic protein-DNA recognition (Levy, Cave, et al., 2000; Shammas, 2017; Ferreiro, et al., 2008; Fuxreiter, et al., 2011). Moreover, the ability to return quantitative affinity measurements for multiple oligonucleotide sequences provides critical information required to probe the mechanistic origins of specificity and extract energetic information from crystallized structures (Farrel, et al., 2016).

The experimental difficulty of quantitatively assaying effects of protein mutations has led to the development of various computational algorithms that use combinations of phylogeny (Ng and Henikoff, 2003; Hopf, et al., 2017; Choi, et al., 2012), structure (Adzhubei, et al., 2010; Blanco, et al., 2018), and prior experimental measurements (Pelossof, et al., 2015) to predict mutational effects on DNA binding. While these computational models have shown some success in predicting if particular disease-associated mutations ablate DNA binding, comparisons between computational predictions and the experimental data generated here establish that prediction accuracies are modest at best and typically underestimate effects of mutations to poorly conserved, non-contacting residues (Miller, et al., 2017; Reeb, et al., 2020). In future work, we anticipate that STAMMP datasets may serve as critical resources for testing, revising, and refining computational algorithms. Furthermore, systematic studies of TFs at this resolution may reveal context dependencies for effects of TF mutations and mechanisms by which sequence variation within structurally related TFs give rise to differences in sequence preference (Berger, et al., 2008; Salisbery, et al., 2011).

Within the cell nucleus, TF-DNA interaction affinities over a wide physiological range precisely specify transcriptional programs (Farley, et al., 2015; Kribelbauer, et al., 2018; Le, et al., 2018), and even subtle decreases or increases to affinity can alter transcription rate and disrupt overlying genetic circuits (Gaudet and Mango, 2002; Meinhardt, et al., 2013; Farley, et al., 2015; Crocker, et al., 2016; Wang, et al., 2017; Bhimsaria, et al., 2018; Le, et al., 2018; Nishizaki, et al., 2020). For most mutations attempting to recapitulate disease-associated human variants, binding affinity was reduced to background levels as previously reported (Wang, et al., 2017). However, one mutant designed to mimic disease-associated orthologs (H255R) increased DNA binding affinity across a broad spectrum of DNA sequences, suggesting that *enhanced* affinity may drive disease by altering transcriptional programs. This observation is consistent with prior studies suggesting TF affinities must be appropriately tuned for function and that very high-affinity DNA binding sites can be statistically significantly underrepresented within genomes (Le, et al., 2018; Bruno, et al., 2019). Moreover, these results imply that in some cases, appropriate therapeutic interventions could attempt to disrupt (rather than restore) binding.

In future work, the ability to express and functionally characterize hundreds of TF mutants in a single experiment at low cost enables a wide variety of precision medicine applications. STAMMP could be used to systematically assess effects of disease-associated TF variants, thereby providing a high-throughput method for assessing functional consequences of variants of unknown significance. STAMMP could also be extended towards testing compounds for their potential to drug transcription factors and serve as new cancer therapeutics (Struntz, et al., 2019; Lambert, M., et al., 2018).

## Acknowledgements

P.M.F. acknowledges the support of an Alfred P. Sloan Foundation fellowship and is a Chan Zuckerberg Biohub Investigator. A.K.A. and N.V.D. acknowledge the support of the National Science Foundation (GRFP). A.K.A. also acknowledges support from the ChEM-H Chemistry/Biology Interface (CBI) Predoctoral Training program. C.J.M. was supported by a Canadian Institutes of Health Research (CIHR) Postdoctoral Fellowship. D.A.M. acknowledges support from the Stanford MSTP program and the Xu Family Foundation Stanford Interdisciplinary Graduate Fellowship (SIGF) affiliated with Stanford ChEM-H. The authors thank Scott Longwell for experimental pipeline assistance and Kara Brower for microfluidic mold fabrication. We thank Connor Horton and Michael Hayes for critical feedback on the manuscript.

## Author Contributions

Conceptualization: A.K.A. and P.M.F.; Formal Analysis: A.K.A. and P.M.F.; Investigation: A.K.A. and N.V.D.; Resources: A.K.A., C.J.M., D.A.M.; Writing – original draft: A.K.A. and P.M.F.; Writing – review & editing: All authors; Supervision & Funding Acquisition: P.M.F.

