## Supplemental methods and figures for "High-throughput binding affinity measurements for mutations spanning a transcription factor-DNA interface reveal affinity and specificity determinants"

**Mold and device fabrication:**

Flow and control molding masters were fabricated as described previously (Fordyce, et al., 2012; Le, et al., 2018). Two-layer microfluidic devices were then cast from these molds using polydimethylsiloxane (PDMS) polymer (RS Hughes, RTV615). Control layers of the microfluidic device (**Fig. S1,** orange) were generated by pouring 60 grams of PDMS (1:5 ratio of cross-linker to base) onto the molds, degassing to remove all air in a vacuum chamber under vacuum for ~45 minutes, and baking for 40 minutes at 80ºC in a convection oven. After this step, control layers for each device were cut out and removed from the wafer and the fluid line inlets were punched using a catheter hole punch (SYNEO, CR0350255N20R4) mounted onto a drill press (Technical Innovations). The flow layer (**Fig. S1**, blue/green) was generated by spin-casting PDMS (1:20 ratio of cross-linker to polymer) onto the molds at 266 rpm for 10 sec, followed by 1750-1850 rpm for 75 seconds. Layers were relaxed on a flat surface for 10 minutes at room temperature prior to baking at 80ºC for 40 minutes in an oven. Cut and punched device control layers were then aligned to flow layers remaining on master molds manually using a stereoscope. Aligned devices were then baked for 40 minutes at 80 ºC in an oven, excised from the molds using a scalpel, and the remaining flow-layer fluidics inlets made using the same catheter punch as above.

**High-throughput QuikChange Mutagenesis:**

Pho4 plasmid: Prior to Pho4 mutant library generation, we generated a version of the PURExpress control expression plasmid (containing a T7 promoter and an ampicillin selectable marker) in which we inserted the full coding sequence for WT Pho4 fused to a C-terminal monomeric GFP tag (Zacharias, et al., 2002) with an intervening gly-ser linker (GGGSGGGGSG). This insertion was sequence-validated via Sanger sequencing and all subsequent mutagenesis used this sequence as a plasmid template.

Primer design: Mutagenic primers were designed using an in-house automated script (https://github.com/FordyceLab/designQuikChangePrimers). Briefly, the program takes as input a list of desired mutations (e.g. “H255R” for His 255 to Arg mutation) and the protein ORF sequence. For each desired mutation, the script generates different primer candidates by varying the primer length and the position of the mutation site relative to the center of the primer. Candidates were scored using heuristics based on the manufacturer’s recommendations included in the QuikChange (Agilent) protocol. First, we selected the mutagenic codon requiring the smallest number of nucleotide mismatches. Next, we calculated the primer scores as follows: (1) we calculated the annealing temperature ($T_m$) according to the formula given in the QuikChange manual and scored primers such that a maximum score of 2 was given for 78≤Tm<82°C and this score linearly decreased (with slope=1) for each 1°C change outside this range; (2) we added an additional score of 1 for a G-C base pair at the 5’ or the 3’ end; (3) we subtracted between 0.25 and 1 if the primer was predicted to have 3’ self-complementarity, depending on the degree of predicted self-complementary; (4) we subtracted 0.25 from the score if the first two 5’ or 3’ bases were the same, to minimize the risk of slippage at the ends; (5) we penalized primer candidates longer than 38 bp (as long as Tm was > 79°C) to reduce oligo synthesis costs; and (6) we calculated predicted primer hairpin temperatures using the primer3-py ‘calcHairpin’ function (Untergasser, et al., 2012) and performed an additional round of optimization if the primer had a predicted hairpin with melting temperature >50°C to decrease this melting temperature, if possible. The final primer was chosen based on the highest cumulative score. Additional details regarding primer optimization calculations are available in the Github repository. Optimized primers were subsequently ordered from IDT (Integrated DNA Technologies) at the 10 nmol synthesis scale in a 96-well plate format with standard desalting purification. These primers were normalized at 6 nmol per well with forward and reverse primers encoding mutations premixed in each well and shipped dry.

QuikChange Mutagenesis: Primers were resuspended in 120 µL of Milli-Q $H_2O$ and left at room temperature to allow primers to solubilize for approximately 1 hour, creating working stock solutions of 50 µM for downstream PCR. Using a 96-channel manual pipettor (Liquidator, Rainin), we added 5 µL of solubilized primers to 195 µL of Milli-Q water to dilute to a working concentration of 1.25 µM and mixed well by pipetting up and down. Finally, we transferred 6 µL of these diluted primers to a new 96-well plate before adding QuikChange reaction Master Mix to all wells (as described below).

We prepared the QuikChange master mix (Agilent Technologies, New England Biolabs) by scaling the single reaction recipe as necessary and keeping on ice prior to use:
*Per single reaction:*
14 µL deionized $H_2O$ (*e.g.* Milli-Q $H_2O$)
2.5 µL 10X Pfu buffer (AD)
0.25 µL Plasmid template (100 ng/µL)
1.25 µL DMSO (5% v/v)
0.5 µL dNTPs (final concentration: 200 µM)
0.5 µL Pfu turbo polymerase (AD) (0.05 Units)
6 µL Forward and Reverse primers (300 nM)

After preparation of the Master Mix, we added 19 µL of this Master Mix into each well using a multichannel pipette and mixed well. We then sealed plates with foil, centrifuged briefly, and placed them into the thermocycler for the PCR reaction according to manufacturer's protocols (Agilent QuikChange Manual).

Next, we treated reactions with Dpn1 enzyme (New England Biolabs, R0176L) to digest any remaining WT plasmid. To do this, we took 10 µL of the above mutagenesis reaction, and added 10 µL of Dpn1 reaction mix (1 µL Dpn1, 2 µL CutSmart Buffer, and 7 µL deionized $H_2O$). These reactions were well-mixed via pipetting and incubated on a thermocycler with the following protocol.

37ºC, 3 hrs

80ºC, 20 min
4ºC, hold

We then used 1 µL of each reaction to transform 5 µL of *E. coli* DH5α cells (New England Biolabs, C2987I). The cells and PCR product were left on ice for 30 minutes, heat-shocked at 42ºC for 30 seconds, and recovered in 300 µL SOC Medium (NEB, B9020S) for 1 hour prior to plating on LB agar plates supplemented with ampicillin (100 µg/mL). Plates were kept at 37ºC overnight to grow colony transformants. Single colonies from these plates were then picked and grown at 37ºC in 6–8 mL of LB medium supplemented with ampicillin (100 µg/mL) overnight; plates were subsequently stored at 4 ºC in case additional colonies need to be picked. We miniprepped plasmids using Qiagen miniprep reagents and validated proper mutagenesis via Sanger sequencing. Colonies were re-picked if the sequencing revealed a wildtype Pho4 clone, an errant mutation elsewhere in the construct, or poor sequencing quality. In cases where no transformants appeared after the mutagenesis reaction, we repeated QuikChange reactions with manually designed primers.

**Plasmid array printing:**
Prior to printing, we transferred 10 µL of the plasmid solution from each well of the 96 well to 2 different wells within 384 well plates (Thermo Scientific, AB-1055) using a Biomek FX Automated Workstation (Beckman Coulter, model A31843), recording plasmid locations to map mutants to chambers in downstream experiments. To standardize well volumes, we then evaporated all wells

within the 384 well plates to dryness and re-solubilized in 12–15 µL of print solution formulated as follows:

1% (10 mg/mL) Bovine Serum Albumin (Sigma Life Science, B4287-25G)
200mM (11.65mg/mL) NaCl (Sigma Life Science, 71376-1KG)
12mg/mL trehalose dihydrate (Sigma Life Science, T9531-25G)

These ingredients were combined and dissolved in Milli-Q H2O and filter sterilized prior to use (Millipore, SE1M179M6). When not in use, we stored the print solution and plates with plasmids at 4ºC. For longer term storage, we sealed plasmid plates with foil and stored at -20ºC.

Prior to printing, all plates were defrosted at 4ºC overnight and then centrifuged at 4ºC (2000g for 10 minutes). We arrayed plasmids onto epoxysilane-coated 2"x3" glass slides (Thermo Scientific, UCSF2X3-C50-20) using a custom built microarrayer outfitted with silicon pins (Parallel Synthesis Technologies, SMT-S75). To prevent contamination of DNA during prints, pins were washed twice in near-boiling water for 10 seconds, followed by vacuum drying for 8 seconds. After arrays dried overnight, we aligned fabricated PDMS devices onto printed plasmid arrays such that each plasmid spot was isolated in its own unique chamber within the device. Devices were then bonded to the glass slides by baking for 4–12 hours at 95ºC on a hotplate (Torrey Pines Scientific) prior to running experiments.

**Preparation of fluorescently-labeled dsDNA for binding assays:**
DNA sequences studied: All DNA sequences were designed with a universal complementary 3' region to allow annealing of a single 5' AlexaFluor-647-conjugated DNA primer to all sequences (calculated $T_m$ of this annealing was 37ºC) (Table 1).

| Name | Full Sequence (5' to 3') |
|---|---|
| CCACGTGA | CAATACACTGTTATC AGAC**C CACGTG A**CGAG CTACTCGTTCGGTTATCCGGCGGTATGAC |
| TCACGTGC | CAATACACTGTTATC AGAC**T CACGTG C**CGAG CTACTCGTTCGGTTATCCGGCGGTATGAC |
| ACACGTGA | CAATACACTGTTATC AGAC**A CACGTG A**CGAG CTACTCGTTCGGTTATCCGGCGGTATGAC |
| GCACGTGC | CAATACACTGTTATC AGAC**G CACGTG C**CGAG CTACTCGTTCGGTTATCCGGCGGTATGAC |
| CAACGTGA | CAATACACTGTTATC AGAC**C AACGTG A**CGAG CTACTCGTTCGGTTATCCGGCGGTATGAC |
| CCGCGTGA | CAATACACTGTTATC AGAC**C CGCGTG A**CGAG CTACTCGTTCGGTTATCCGGCGGTATGAC |
| CCATGTGA | CAATACACTGTTATC AGAC**C CATGTG A**CGAG CTACTCGTTCGGTTATCCGGCGGTATGAC |
| CCACGCGA | CAATACACTGTTATC AGAC**C CACGCG A**CGAG CTACTCGTTCGGTTATCCGGCGGTATGAC |
| CCACGTTA | CAATACACTGTTATC AGAC**C CACGTT A**CGAG CTACTCGTTCGGTTATCCGGCGGTATGAC |
| "Universal" | AlexaFluor-647-5'-GTCATACCGCCGGA-3' |

**Table 1.** DNA oligonucleotide sequences used in these experiments. For each sequence, the 8 bp TF consensus site is shown in bold, the single nucleotide variant is highlighted in red, and the universal 3' sequence used for annealing a 5' Alexa-647-conjugated sequence is shown in blue.

Double-stranded DNA preparation and dilution: We ordered nearly all DNA sequences as a single strand from Integrated DNA Technologies (IDT) with standard desalting purification and

‘LabReady’ formulation (100 µM in IDTE buffer, pH = 8.0); for several oligonucleotides ordered dry, we resuspended the oligonucleotide in Milli-Q $H_2O$ to a final concentration of 100 µM (confirmed using a DeNovix instrument). We then converted these ssDNA sequences to fluorescently-labeled double-stranded DNA via: (1) annealing of a universal 5’-AlexaFluor-647-labeled primer to all sequences, and (2) extension using Klenow fragment, exo-. To minimize variation in measured fluorescence intensities between runs, we prepared fresh labeled dsDNA on the day of each experiment.

To begin annealing and extension reactions, we defrosted NEBuffer 2 (New England Biolabs, B7002S) and dNTPs (100 mM, Thermo Scientific) and kept them on ice. Next, we prepared two annealing reactions, formulated as follows:

12 µL single-stranded DNA (100mM)
12 µL “Universal” primer (100uM)
12 µL dNTP mixture (4 mM)
4 µL NEBuffer 2 (10x stock)

We then performed the annealing reaction using the following protocol on a thermocycler:

94ºC, 3 min
Cool to 37ºC over 45 minutes

To extend the annealed universal primer, we removed the tubes from the thermocycler, spun them down using a centrifuge, added the following, and mixed well via pipetting:

8 µL of Milli-Q $H_2O$
1 µL of NEBuffer 2 (10x stock)
1 µL of Klenow enzyme (made in-house)

The tubes were then placed back in the thermocycler for the extension step of the protocol:

37ºC, 60 min
80ºC, 20 min
10ºC, hold

After the extension step, we again centrifuged the tubes using a table-top microcentrifuge and placed them on ice. To remove any aggregates that could clog the microfluidic channels on the device, we sterile filtered reactions with a 0.45um filter spin column (Merck Millipore, UFC30HVNB). Finally, we equilibrated dsDNA reactions in the final assay buffer (10mM Tris-HCl, 100mM NaCl, 1mM DTT, pH 7.5; aliquoted and filtered using 0.45 µM Steriflip vacuum (Millipore, SE1M179M6) using 10K filter spin concentrator columns (Amicon Ultra, UFC501096). To do this, we added 100 µL of the duplexing reaction in the filter spin concentrator, added 200 µL of assay buffer (~300 µL total volume), and spun down to concentrate back to 100 µL via centrifugation (8000-9000g for 8 minutes); this process was repeated 5 times. After the final step, we eluted the DNA by inverting the filter into the collection tube and centrifuging at 3000g for 5 minutes.

For each DNA concentration series, we serially diluted this eluent 1:2 in assay buffer supplemented with 50 µg/mL of UltraPure BSA (ThermoFisher, AM2618) to yield effective final concentrations of ~5 µM, ~2.5 µM, ~600nM, ~300 nM, ~160nM, ~90 nM. For DNA sequences containing mutations in the core binding site, we modified this concentration series to include slightly higher maximum DNA concentrations (~ 7.5 µM, ~ 4 µM, ~ 2 µM, ~ 400 nM, ~ 200 nM, ~40 nM). We noted that addition of UltraPure BSA did not result in changes in affinity. We then calibrated fluorescence intensity to effective DNA concentration by using a DeNovix instrument to measure the absorbance at 260 nm.

**Microscopy instrumentation:**

We performed all measurements on a Nikon Ti-S Microscope with a motorized XY stage (Applied Scientific Instrumentation, MS-2000 XYZ stage), cMOS camera (Oxford Instruments, Andor Zyla 4.2 CMOS), and solid-state light source (Lumencor, SOLA SE Light Engine). We programmed the microscope to perform a gridded acquisition of the device using MicroManager (Edelstein, et al., 2014). This grid was set using a 10% overlap between imaging fields for image stitching (described in Analysis), and all images were collected using a 4X objective lens using a 2x2 bin setting.

**On-chip surface patterning:**

Device connections for reagent introduction: To introduce reagents onto the microfluidic device, reagents were loaded via syringe suction into Tygon tubing (Saint-Gobain, AAD04103) connected to a syringe at one end using a 23-gauge luer connector (McMaster-Carr, 75165A684) and fitted with a blunt-ended steel pin at the other (0.013 in ID x 0.025 in OD x 0.5 in length, New England Small Tube Corporation, NE-1310-02). After loading reagents into the syringe, we inserted the blunt-ended steel pin into the appropriate device port, removed the other end of the Tygon from the Luer-fitted syringe (Brower, et al., 2018), and connected the tubing directly to a custom pneumatic manifold that drives fluid flow via positive pressure (Brower, et al., 2018). All pneumatic control valves were actuated using a custom automated pneumatic control manifold (Brower, et al., 2018) controlled using an in-house python software package (https://github.com/FordyceLab/RunPack).

Surface patterning: Device surfaces were functionalized with surface-immobilized antibody largely as previously described (Aditham, et al., 2018; Le, et al., 2018); however, we included a few modifications. First, valve lines controlling the “button”, “sandwich”, and “neck” valves (**Fig. S1**) were pressurized with 550mM NaCl in Milli-Q $H_2O$ to prevent premature solubilization of the DNA spots by osmotic balancing of fluids between the pneumatic control and reagent flow channels. All other control lines were pressurized with Milli-Q $H_2O$. Second, we controlled all pneumatic valves on the device at a pressure ranging from 35-37 psi and introduced reagents at pressures of 3.5-4 psi.

To begin antibody patterning, we first dead-end filled all device control lines. Next, we flowed biotinylated BSA (2 mg/mL, ThermoFisher Pierce, 29130) to expel all air from device flow layers by opening the inlet and outlet valves; to expel any remaining air bubbles from the flow channels, we closed the outlet valve after 2-3 minutes and continued to apply pressure to dead-end fill the device for an additional 5-10 minutes. After all air bubbles were expelled, we opened the outlet valve to allow reagent flow, and introduced 2 mg/mL biotinylated BSA for an additional 5 minutes with the “button” valves pressurized and then for 30 minutes with the “button” valves open. Next, we flushed the device with phosphate buffered saline (10X stock, Corning, 46-013-CM; diluted to 1X in Milli-Q $H_2O$) for 10 minutes. We then introduced neutravidin (1 mg/mL, Thermo Scientific, 31000) for 30 minutes with “button” valves opened, followed by another PBS wash for 10 minutes. To passivate all device surfaces except those protected by the “button” valves, we the pressurized the “button” valves and introduced biotinylated BSA again for 30 minutes, thereby coating device surfaces and preventing non-specific antibody binding. After an additional 10 minute PBS wash, we introduced biotinylated anti-GFP antibody (100 ug/mL, Abcam, ab6658) into the device for 2 minutes with the “button” valves pressurized (to ensure antibody was evenly distributed throughout the device) and then opened “button” valves and flowed for an additional 13 minutes and 20 seconds, thereby specifically recruiting biotinylated anti-GFP antibodies to exposed neutravidin-coated surfaces beneath “button” valves **(Fig. S2)**. Finally, we washed the device with PBS for 10 minutes. Upon conclusion of these steps, we stored the device with the "button” valves pressurized, the outlet valve shut and the PBS inlet valve open. A light PBS flow kept the device from drying out; we typically completed surface chemistry the night before an experimental assay.

**On-chip TF expression and purification:**

On-chip TF expression: After surface patterning, we expressed all TF variants on the device using the PURExpress *in vitro* transcription/translation system (New England Biolabs, E6800L). First, we combined PURExpress components A (10 µL) & B (7.5 µL) off-chip, mixed gently per the manufacturer's instructions via pipetting, and stored the reaction on ice for 45 minutes, which we observed increased expression yields. After this, we added 1.5 µL of RNAsin (Promega, N2515) and Milli-Q water or DNAse-free water (Promega) to 25 µL. A single 25 µL PURExpress reaction was sufficient for expression on a single device; for multiple devices, we scaled this reaction master mix as necessary.

After incubation, we flowed PURExpress through flow channels for 10 minutes with the "neck" and "button" valves closed and the "sandwich" valve open to completely fill all channels. After this period, we closed the outlet valve, opened the "neck" valves, and closed the "sandwich" and "button" valves to push PURExpress into the plasmid compartments. To ensure that each plasmid compartment was fully filled with PURExpress, we again opened the "button" and "sandwich" valves with the outlet valve closed and continued flowing PURExpress for an additional 2-3 minutes. To begin protein expression, we then opened the "neck" valves (to allow protein expression in the entire chamber), closed the "sandwich" valves and "button" valves (thereby isolating adjacent chambers from one another and protecting the antibody-coated surface), and placed the device on a preheated hot plate (Torrey Pines Scientific) at 37ºC for 45 minutes. To allow GFP to fold and mature, we then removed the device from the hot plate and incubated at room temperature for an additional 2 hours. Finally, we opened the "button" valves to allow expressed GFP-tagged TFs to bind patterned anti-GFP antibodies on the slide surface beneath the "button" valves. During this step, we mounted the device on a Nikon Ti-S microscope (described above) and periodically imaged using gridded acquisition imaging software to monitor build-up of GFP intensity over time.

On-chip TF purification: After allowing TF binding to proceed for 2 hours, we again closed the "neck" valve (to sequester the plasmid compartment from the reaction compartment), closed the "button" valve (to protect surface-immobilized TFs from flow-induced shear, opened the "sandwich" valves, and washed with PBS for 10 minutes to wash away any non-specifically bound TF proteins, thereby purifying TFs within each chamber. To remove TF proteins non-specifically adsorbed to device walls, we additionally washed with TrypLE (1X stock, ThermoFisher, 12604-013) for 15 minutes, washed with 2 mg/mL biotinylated BSA for 15 minutes to re-passify device walls, and finally washed with PBS for 10 minutes. After this step, we removed all reagent lines except for PBS and washed the inlet manifold with PBS for 10 minutes to remove trace amounts of trypsin that could otherwise damage proteins during the assay.

**DNA binding measurements.**

After on-chip recombinant protein expression and purification, we prepared the device for DNA incubation. Briefly, we replaced the fluidics line containing PBS with a new line containing assay buffer and again washed the inlet manifold of the device for 10 minutes. To equilibrate expressed TFs with assay buffer, we flowed assay buffer across the device for 5 minutes with the "button" valves closed, then opened the "button" valves and flowed for an additional 2 minutes; this process was repeated twice. We followed this with another 50-minute buffer equilibration period, in which "button" valves were open, exposing TFs to assay buffer. During this preliminary equilibration step, we connected Tygon tubing containing each labeled dsDNA concentration to be assayed into the device, allowing all subsequent experimental steps to be automated using custom in-house software (https://github.com/FordyceLab/RunPack).

For each concentration of DNA (from the lowest to the highest), we: (1) closed "neck" and "button" valves and opened "sandwich" and inlet and outlet valves; (2) flowed labeled DNA across the device for 10 minutes; (3) closed "sandwich", inlet, and outlet valves; (4) opened "button" valves (to allow surface-immobilized TFs to interact with soluble DNA); (5) incubated for 50 minutes (to allow reactions to come to equilibrium); (6) imaged all chambers within the device in the Cy5 (DNA) channel (to quantify intensities and calculate the concentration of soluble DNA available for binding in each chamber); (7) closed "button" valves (to trap TF-bound DNA); (8) washed with assay buffer for 10 minutes; and (9) imaged all chambers in both the GFP (TF) and Cy5 (DNA) channels to quantify the relative intensities of trapped species in each chamber. To quantify free DNA in solution at each step, we imaged the device in the Cy5 channel at exposures ranging from 30 ms to 150 ms to ensure an adequate dynamic range for downstream analysis. To quantify surface-immobilized TF intensities and intensities of bound DNA, we acquired 500 ms exposure images in the GFP channel and 3000 ms exposure images in the Cy5 channel. To image the entire device, we tiled acquisitions across the device (typically 7 x 7 gridded images).

**Data Analysis:**
We performed all analysis in Python, with exceptions as noted below.

Image processing and quantitation: To correct for position-dependent variation in excitation illumination and collection efficiency, we applied a flatfield correction to all GFP and Cy5 fluorescence images using correction images as previously described (Thorn, 2014). We also collected initial images of devices prior to the start of each experiment for every channel and exposure time to collect background measurements to calculate optimal flatfield correction parameters. Tiled acquisition images were then stitched to generate a single large image of each device using an in-house image stitching program (https://github.com/FordyceLab/ImageStitcher).

Each DNA concentration measurement yielded 3 stitched images: (1) a "Prewash Cy5" image in the Cy5 channel used to calculate the concentration of free DNA available for binding; (2) a "Postwash GFP" images used to quantify the amount of surface-immobilized TFs at each step of the binding assay; and (3) a "Postwash Cy5" image used to quantify the amount of fluorescently-labeled DNA bound to surface-immobilized TFs at each step.

To quantify Alexa-647-labeled DNA intensities within each chamber (for the "Prewash Cy5" image) and labeled DNA and TF intensities beneath the "button" valve (for the "Postwash GFP" and "Postwash Cy5" images), we used a custom, in-house python image processing package (https://github.com/FordyceLab/ProcessingPack). Briefly, we first identified the four corners of the device by marking the centers of the top left, top right, bottom left, and bottom right corner binding reaction chambers using the "Prewash Cy5" image associated with a high introduced DNA concentration. The other chamber coordinates were initially approximated by using these corners as vertices of a 28-column x 56-row grid (chamber dimensions of the PC1k device); we then employed a Hough Transform to find chamber centers. We then used these same coordinates for quantify median fluorescence chamber intensities for all chambers across all 6 measured DNA concentrations. Finally, median chamber intensities were converted to DNA chamber concentrations using per-chamber DNA calibration curves (as described below).

To quantify fluorescence intensities associated with surface-immobilized TFs and trapped bound DNA, we first identified "button" centroid locations and perimeters using the first "Postwash GFP" image as a reference. "Button" centroid positions were approximated using a grid search to maximize the fluorescence intensity within a circular area slightly larger than the expected physical size of the buttons. We then applied these same feature locations to all "Postwash GFP" and "Postwash Cy5" images, indexed by the DNA concentration step at which they were acquired. To quantify "button"-associated GFP and Alexa-647 intensities within each chamber, we summed the total fluorescence spot intensity in each channel and then subtracted contributions from local background (calculated by summing the intensities of an annulus surrounding the spot area and then normalizing this value by the relative ratio of the "button" and annulus areas). Finally, we associated all chamber and "button" intensities with a particular mutant using a custom Python script that maps plate well location (and associated mutant ID) to reaction chambers within the device.

Quality control and binding curve generation: Prior to generating binding curves, we employed several quality control checks to the raw imaging data: (1) we computed a linear regression between observed intensities for each chamber in the "Prewash Cy5" image and the concentration of introduced DNA, inspected $r^2$ values associated with each fit (**Fig. S3**); (2) we compared the distributions of GFP intensities between plasmid-containing chambers and empty chambers and eliminated chambers with intensities below an experiment-specific threshold ($1.5x10^6$ - $2.5x10^6$ RFU); (3) we eliminated any chambers with measured "Postwash Cy5" median intensities below background levels; and (4) we manually examined raw images to eliminate chambers containing any unusual particulates or debris leading to aberrant DNA binding curves.

To generate per-chamber binding curves, we then plotted the measured ratio of intensities beneath the "button" valve (Alexa-647-DNA/TF-GFP from "Postwash Cy5" and "Postwash GFP" images) against the DNA concentration at each assay step.

Calculating $K_d$ values: To determine measured $K_d$ values for each chamber, we globally fit concentration-dependent binding curves to a single-site binding model:

$$R = \frac{R_{max} \cdot [DNA]}{K_d + [DNA]}$$

In this equation, $R_{max}$ represents the ratio of DNA/TF fluorescence at saturating DNA concentrations, and $K_d$ represents the DNA concentration at which half of the surface-immobilized TFs are occupied by DNA. To enhance the accuracy of estimated $K_d$ values even for oligonucleotides with binding affinities above the highest measured DNA concentration, we initially determined a global $R_{max}$ value for all mutants by calculating the median Alexa-647-DNA/TF-GFP ratio for the top 10% of DNA/protein measurements at the final DNA concentration within the assay. We then fit all curves using this constant $R_{max}$ across all chambers within a given experiment, optimizing only the per-chamber $K_d$ value via nonlinear least squares fitting. Over several experiments for experiments containing only flanking nucleotide mutations (where many mutants reached saturation at the maximum [DNA]), this $R_{max}$ value was largely consistent (0.67 ± 0.012, median ± SEM) (**Fig. S13**). For experiments assessing binding to oligonucleotides containing mutations in the core consensus site, this median $R_{max}$ value was typically lower, and so all curves were fitted using $R_{max}$ values of 0.66 or 0.68. All binding curves and all returned $K_d$s for all chambers across all experiments are available as Supplemental Files.

To calculate ΔΔGs across a given experiment, we used the following formula:

$$\Delta\Delta G = RT \cdot \ln\left(\frac{K_d}{K_{d,ref}}\right)$$

where R is the gas constant ($1.987 \cdot 10^{-3}$ kcal/(K · mol)), T is 298 K, and $K_{d,\ ref}$ is the median $K_d$ measurement for the wildtype PHO4 protein interacting with the DNA oligonucleotide sequence used in a given experiment.

Determining statistical significance from wildtype Pho4 & from background: To identify mutants with statistically significant differences in DNA binding from the WT protein, we compared the distribution of all measured ΔΔG values for every mutant against the distribution of all measured ΔΔG values for the WT Pho4 protein using an independent, two-tailed T-test (assuming unpooled variance). We used a Bonferroni corrected p-value assuming a normal *p*-value of 0.05 and 213 measured mutants ($p < 0.0003$) as a threshold of significance.

To identify mutants with binding that was statistically significantly different from background (nonspecific) DNA binding on the assay, we compared the distribution of all measured ΔΔG values for every mutant against the distribution of all measured ΔΔG values for an A299D mutant that lacks DNA binding and again used a Bonferroni correction to determine a conservative threshold for significance. To select A299D as an appropriate mutant to use, we compared the Cy5 signal intensities for empty chambers (without expressed TFs) with those of several TF variants at the highest DNA concentration assayed for two sample experiments. Measured fluorescence intensities for two null-binding mutants, H257P and A299D, were consistently similar to those of the empty chambers (**Fig. S7**). As A299 is not involved in DNA binding or close to DNA binding residues, we selected A299D for comparison.

**Predicting likely effects from phylogeny:**
Analysis of Pho4 variants via PROVEAN: To attempt to predict whether certain Pho4 mutants would be 'damaging' for function, we used the PROVEAN (PROtein Variation Effect Analyzer) software tool (Choi, et al., 2010; Choi, et al., 2015) as provided using default cutoff score of -2.5.

To generate a receiver operating characteristic (ROC) curve, we compared the PROVEAN scores to the ΔΔG values for the reference DNA sequence 5'-CCACGTGA-3'. We considered mutants that were between -0.5 kcal/mol and 1 kcal/mol (inclusive) as indistinguishable from wildtype (ie. a benign variant). For ROC analysis, we varied the prediction cutoff score -10 to 3 in increments of 0.5 and calculated four quantities at each cutoff:

(1) true-neutral (TN): statistically indistinguishable binding from wildtype and predicted benign;
(2) false-neutral (FN): statistically significantly different binding from wildtype Pho4 and predicted neutral;
(3) true-deleterious (TD): distinguishable from wildtype Pho4 and predicted deleterious; and
(4) false-deleterious (FD): indistinguishable from wildtype and predicted deleterious.

At each threshold, we calculated true positive rate (TP) and false positive rate (FP) using the following formulae: TP = TN/(TP+FD) and FP = FN/(FN+TD). We then calculated the area under the curve (AUC) for the ROC using the trapz function in Python (NumPy). As a negative control, we also calculated ROC and AUC values for 25 trials in which p-values were scrambled.

Entropy Calculations: We calculated entropy at every Pho4 position using a publicly available deposited sequence alignment for bHLH proteins (bHLH Family identifier: PF00010) We then trimmed columns in the multiple sequence alignment to include non-gapped positions in the Pho4 reference sequence. Entropy was calculated using the information_content method within the Biopython package with the expected frequency of each residue at every position within Pho4 at the default setting for the Biopython information_content method.

**Identifying 'affinity' and 'specificity' mutants:**
Affinity *vs.* specificity analysis: To test if mutations globally enhanced affinity or altered specificity, we considered all data from oligonucleotides containing 'flanking' nucleotide mutations (5'-CCACGTGA-3', 5'-GCACGTGC-3', 5'-ACACGTGA-3', and 5'-TCACGTGC-3') *and* 'core' nucleotide mutations (5'-CCACGTGA-3', 5'-CAACGTGA-3', 5'-CCGCGTGA-3', 5'-CCATGTGA-3', 5'-CCACGCGA-3', and 5'-CCACGTTA-3'). Because their affinities were increased beyond assay resolution, we excluded A289K and A289R from analysis for all 'flanking' nucleotide mutations. For the DNA sequence 5'-TCACGTGC-3', we also noted that H255R raised affinity beyond assay resolution and, for this DNA sequence, also excluded this mutant. For each mutant and for each oligonucleotide sequence, we then calculated the normalized change in affinity for that mutant relative to all Pho4 constructs ($K_{d,mutant}/K_{d,\text{median for all mutants}}$). For each mutant, we then calculated the median and standard deviation values of this metric to identify constructs that

consistently increased or decreased affinity to all oligonucleotides ('affinity mutants') or had a particularly large variance in values ('specificity mutants').

Affinity-enhancing mutants: We then identified 'affinity-enhancing' mutants using a T-test against WT Pho4 using a Bonferroni-corrected *p*-value as the significance threshold and considering pools of 'flanking' and 'core' mutant oligonucleotides separately (**Fig. S16**). To arrive at a final list, we further required that mutants enhance affinity by at least 0.3 kcal/mol (*i.e.* >2 standard deviations for WT Pho4 $\Delta\Delta G$ measurements for oligonucleotides containing 'flanking' mutations (median $\Delta\Delta G$ ~ 0 kcal/mol; standard deviation: 0.13 kcal/mol)) for *either* 'flanking' or 'core' mutations. This allowed, for example, consideration of mutants for which binding affinity was increased beyond reliable measurement for flanking nucleotide mutations (eg. A289R, A289K) but was measurable for core site mutations. We then categorized residues based on whether they were within or outside the DNA binding domain, resulted in altered charge of the TF, was at a solvent-facing residue, or was another type of mutation.

Identifying mutants with 'altered' yet detectable DNA binding: To identify mutations that altered DNA binding relative to WT, we again aggregated affinity measurements for all oligonucleotides and grouped measurements by protein mutant. For each mutant and for each oligonucleotide sequence, we then calculated the change in binding affinity relative to the affinity of the WT Pho4 construct for that oligonucleotide sequence:

$$\Delta\Delta G_{\text{mutant,oligonucleotide}} = RT \cdot \ln\left(\frac{K_d}{K_{d,ref}}\right)$$

We then performed a T-test comparing the distribution of $\Delta\Delta G$ values for each mutant relative to the distribution of $\Delta\Delta G$ values for the WT Pho4 construct. Using a Bonferroni-corrected *p*-value threshold ($p < 0.05/215$ mutants), we identified 163/215 mutants with statistically significantly different binding from the WT Pho4 for at least 1 oligonucleotide. Next, we tested whether binding was statistically significantly different from background binding by comparing the distribution of $\Delta\Delta G$ values for each mutant with that of the DNA binding-deficient A299D mutant. Using the same *p*-value threshold, we identified 184/215 mutants that retained detectable binding across at least one oligonucleotide. Finally, we considered the union of these 2 sets (133 mutants) to generate a list of mutants that statistically significantly altered but did not ablate DNA binding.

Identifying combinations of residue and nucleotide mutations that preserved physiological binding: To identify particular residue/nucleotide mutations that altered but preserved physiologically-relevant levels of binding, we: (1) considered the aggregate dataset of 1849 $K_d$ measurements for all mutants and oligonucleotides, (2) identified mutants with binding that was statistically significantly different from WT but still detectable (as above), (3) filtered out any measurements with $K_d$ values *higher* than the median Kd measured for WT Pho4 interacting with the known physiologically relevant low affinity 5'-C CACGTT A-3' site (*e.g.* mutant/nucleotide combinations with weaker binding) ($K_d = 12 \pm 0.2$ μM), and (4) filtered out TF mutations at known DNA nucleotide-contacting residues (*i.e.* Pho4 positions 252, 255, 259, and 263). Finally, we quantified the fraction of combinations of these TF and DNA mutations that altered yet preserved binding that involved a 'flanking' nucleotide mutation. Through this analysis, we found: (1) 1063/1849 combinations of TF and DNA mutations yielded altered binding affinities that were tighter than that measured for WT Pho4 interacting with the 5'-C CACGT**T** A-3' sequence, (2) that 648/1849 of these combinations were also statistically significantly different from WT, and (3) that 435/648 of these combinations involved 'flanking' nucleotide substitutions (67%).

**Helical propensity analysis:**

Predicted energetic changes resulting from mutation-dependent alterations in helical propensity were taken from prior measurements (O'Neil, KT, DeGrado, WF. *Science*, 1990., Table 1 of publication). To determine the degree to which changes in measured DNA binding affinity could be explained by predicted changes in helical propensity, we plotted measured differences in DBA binding affinity (ΔΔG for Pho4 mutant variants relative to WT Pho4 for the consensus DNA sequence) *vs.* predicted ΔΔG values from changes in helical propensity for the substituted *vs.* the native residue at each position and performed a linear regression. We performed this comparison for solvent-exposed mutations within helix 1 (where we expect changes in helical propensity to dominate observed energetic effects) **(Fig. 5C)**, at nucleotide-contacting residues (where we expect changes in helical propensity to have an effect but that overall energetic changes are dominated by altered hydrogen bonding) **(Fig. S17A)**, and within the loop region (where expect changes in helical propensity to have no correlation with changes in measured binding) **(Fig. S17B)**. For mutations within the loop region, we excluded backbone-contacting residues and two proline residues likely to have additional effects on binding affinity.

**Double mutant cycle analysis:**

Comparing differential effects of TF mutations: Identifying epistasis across the TF-DNA interface requires 4 affinity measurements: (1) WT Pho4 binding a 'reference' DNA sequence, (2) mutant Pho4 binding a 'reference' DNA sequence, (3) mutant Pho4 binding a 'reference' DNA sequence, and (4) mutant Pho4 binding a 'mutant' DNA sequence. Prior to double-mutant cycle analysis, we: (1) eliminated any TF-DNA combinations with < 4 replicate measurements, (2) excluded measurements to TF mutants due to poor fitting quality for the reference DNA sequence (described above in "Affinity *vs.* specificity analysis"), and (3) $\log_{10}$-transformed all data.

For each mutant cycle, we then: (1) identified TF/DNA mutant combinations with 'detectable' binding (assessed using a T-test comparing measured affinities with those obtained for the binding-deficient A299D mutant interacting with the same oligonucleotide), (2) eliminated TF mutants lacking detectable binding for *both* oligonucleotides (retaining TF mutants with detectable binding for at least one oligonucleotide), (3) plotted measured log-transformed affinities for each mutant (median ± SEM) for the 'mutant' DNA sequence *vs.* the 'reference' DNA sequence (**Fig. S18**), (4) calculated a linear regression of these pairwise comparison values, (4) calculated residuals for each mutant relative to this linear regression, (5) computed the Z-score of the residual (ie. standardized residual) for each mutant relative to all mutants ($Z_i = r_i/SD$, where $r_i$ is the residual for mutant *i*, and SD is the standard deviation of the total distribution of residuals), and (6) identified all mutants with $|Z_i| \geq 2$ for further investigation. For each of these candidate mutants, we then directly compared observed binding curves with those predicted if the measured mutational effects on affinity were purely 'additive':

$$K_d^{TF_{mut},\, DNA_{mut}} = \frac{K_d^{TF_{mut},\, DNA_{WT}} \cdot K_d^{TF_{WT},\, DNA_{mut}}}{K_d^{TF_{WT},\, DNA_{WT}}}$$

Combinations of TF and DNA mutations were considered epistatic if raw binding curves clearly deviated from concentration-dependent binding predicted by this 'additive' $K_d$.

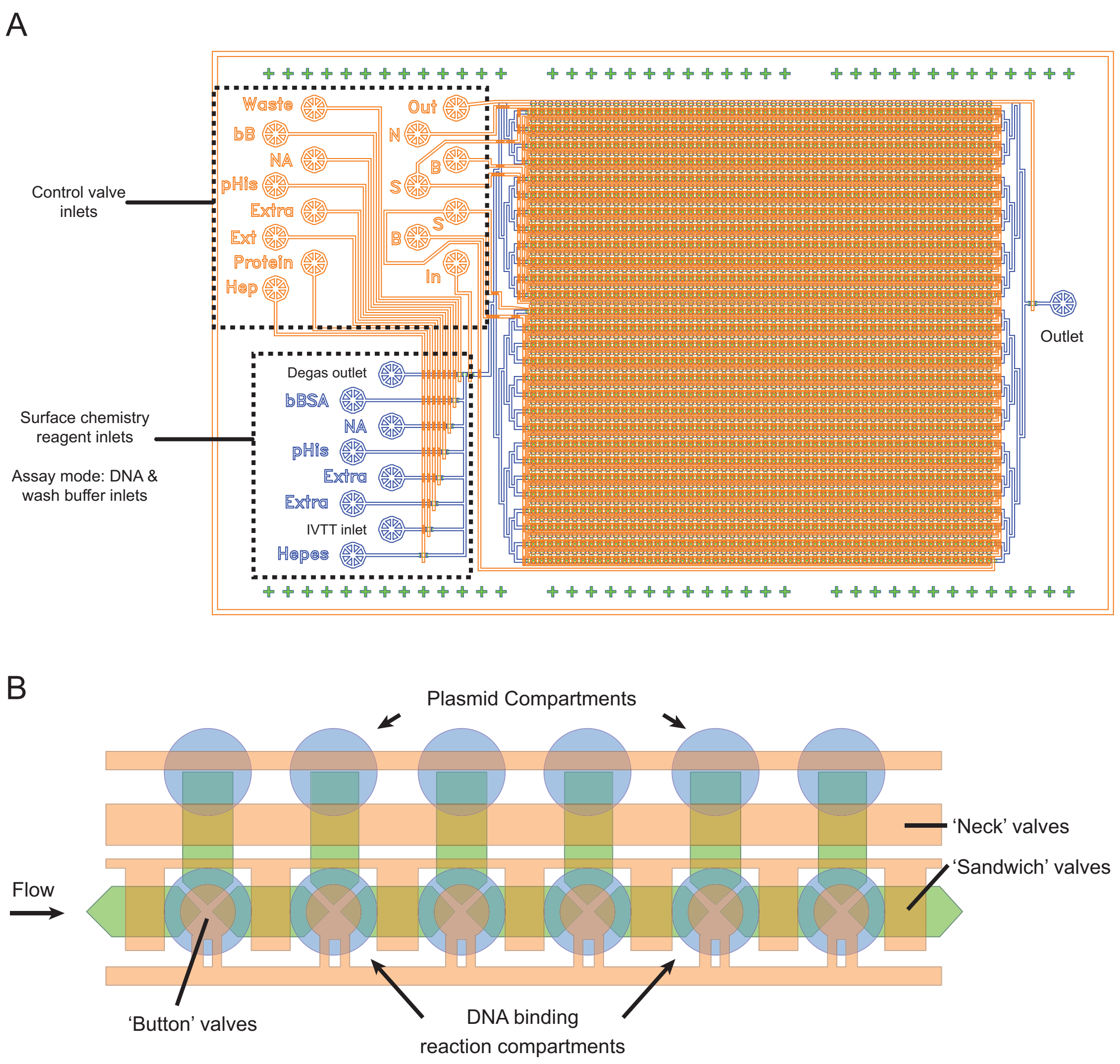


**Figure S1 (Related to Figure 1).** Architecture of MITOMI microfluidic device. **(A)** Detailed diagram of MITOMI microfluidic device with valve inlets and outlets annotated showing reagent flow channels (blue) and pneumatic valve channels (orange) that control flow of reagents in device. **(B)** Magnified view of device chambers showing plasmid and DNA compartments and "neck," "sandwich," and "button" valves.

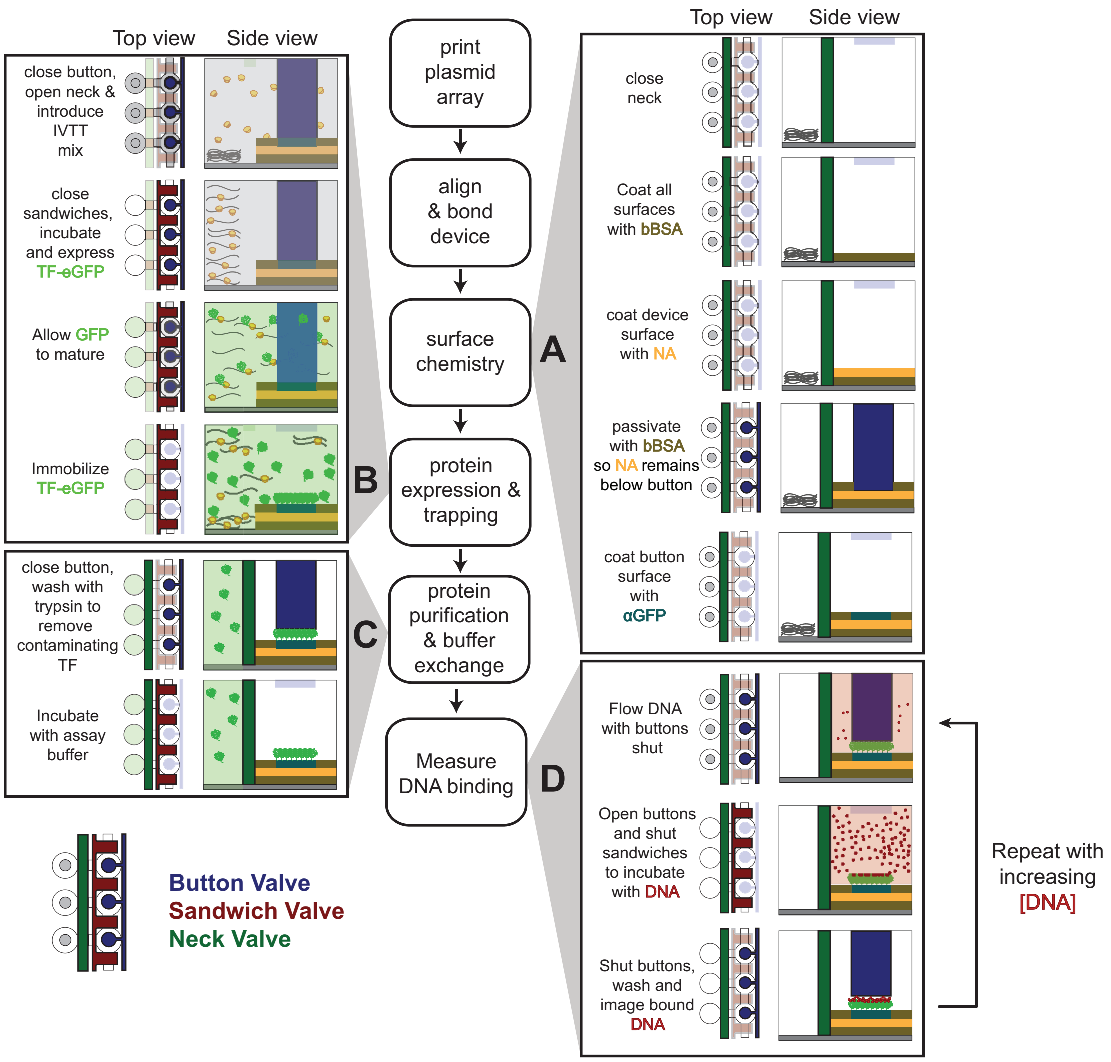


**Figure S2 (Related to Figure 1).** Detailed experimental workflow. **(A)** Surface chemistry to pattern antibody in device for subsequent TF immobilization. **(B)** Expression of GFP-tagged TFs via *in vitro* transcription/translation with sandwich valves depressed to prevent cross contamination. After allowing GFP to fold and mature, button valves are opened to bind expressed protein to patterned antibody-coated surfaces. **(C)** Trypsin digest of nonspecifically bound TFs and equilibration of trapped TF in assay buffer. **(D)** Schematic showing introduction of DNA at a single concentration to quantify binding, with measurements repeated over multiple DNA concentrations to generate a concentration-dependent binding curve.

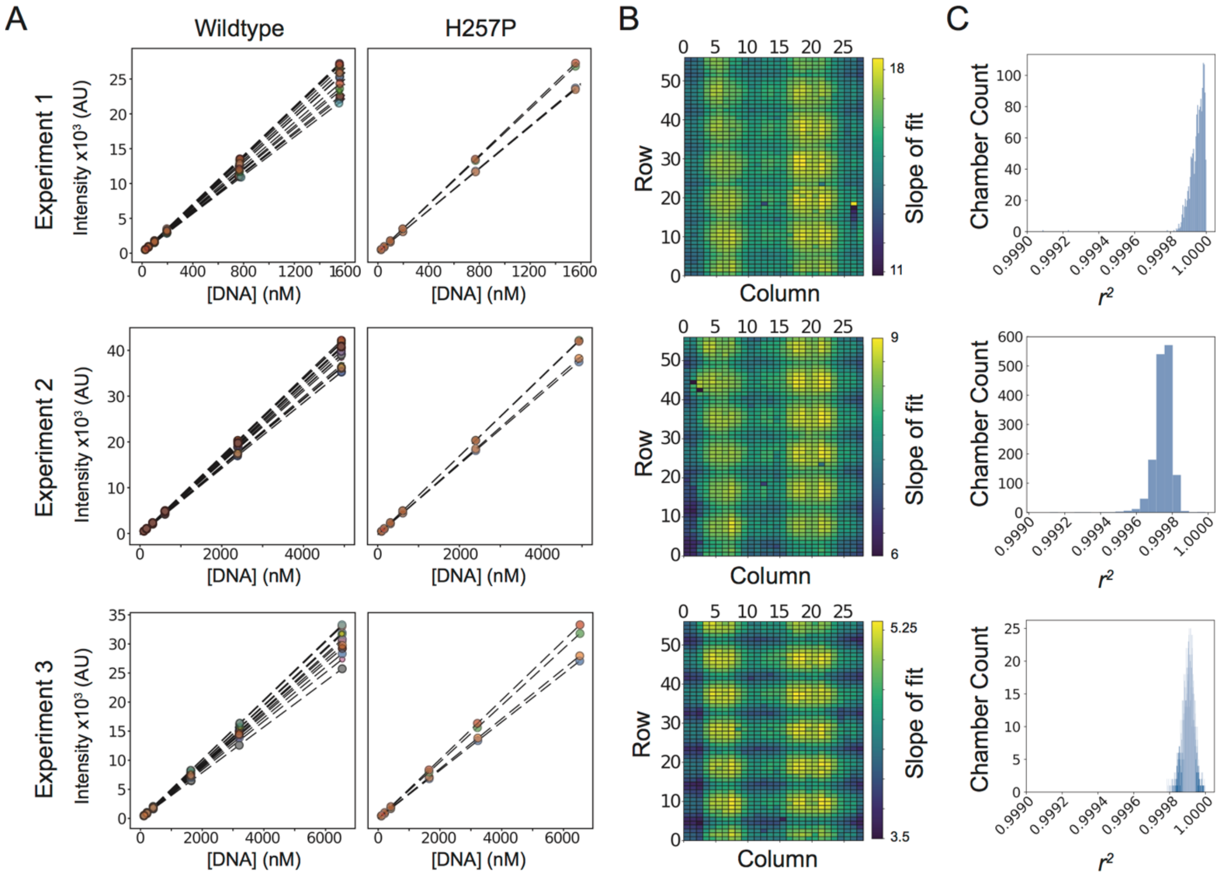


**Figure S3 (Related to Figure 1).** Calibration curves relating fluorescence intensity and DNA concentration. **(A)** Chamber calibration curves for representative sample chambers containing 2 Pho4 constructs (WT and H257P) across 3 experiments showing individual points (colored by device chamber) and associated linear fits (black dashed lines). **(B)** Heatmaps showing linear fit slope as a function of chamber position within device. Slopes vary by approximately 2-fold across the device, with lowest slopes corresponding to outer edges of the microscope field-of-view. **(C)** Goodness-of-fit distributions for calibration curves.

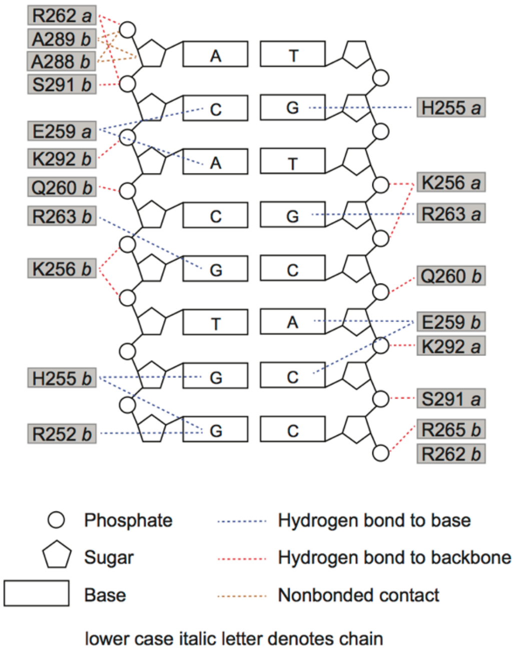


**Figure S4 (Related to Figure 2).** Amino acid residue and DNA contact map of Pho4 (adapted from Shimizu, et al., 1997).

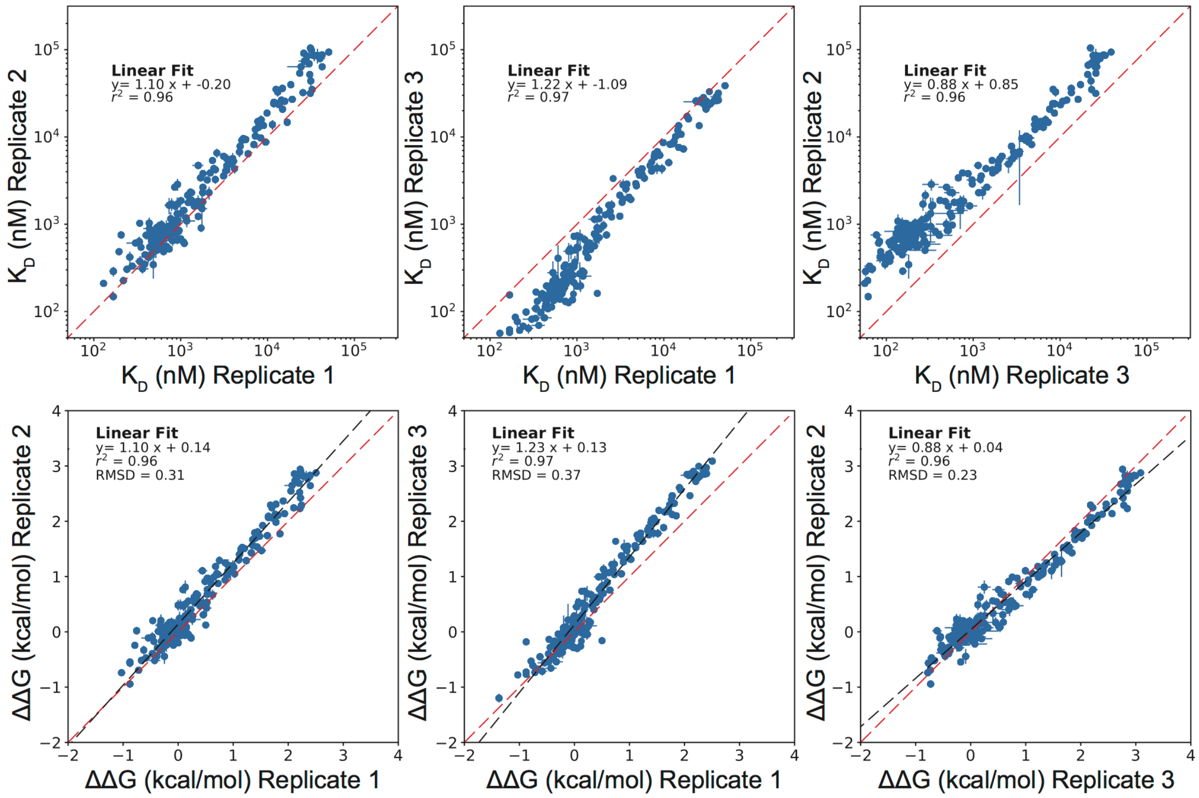


**Figure S5 (Related to Figure 3).** Pairwise comparison of per-mutant $K_d$s (top row) and ΔΔGs (bottom row) for all TF mutants across 3 experiments for reference DNA sequence 5'-C CACGTG A-3'. Points indicate median affinities (± SEM) for each TF mutant. ΔΔGs reflect relative differences in binding energy relative to wildtype and vary by <0.4kcal/mol across experiments. Linear fits are indicated by black dashed lines; identity lines are indicated by red dashed lines.

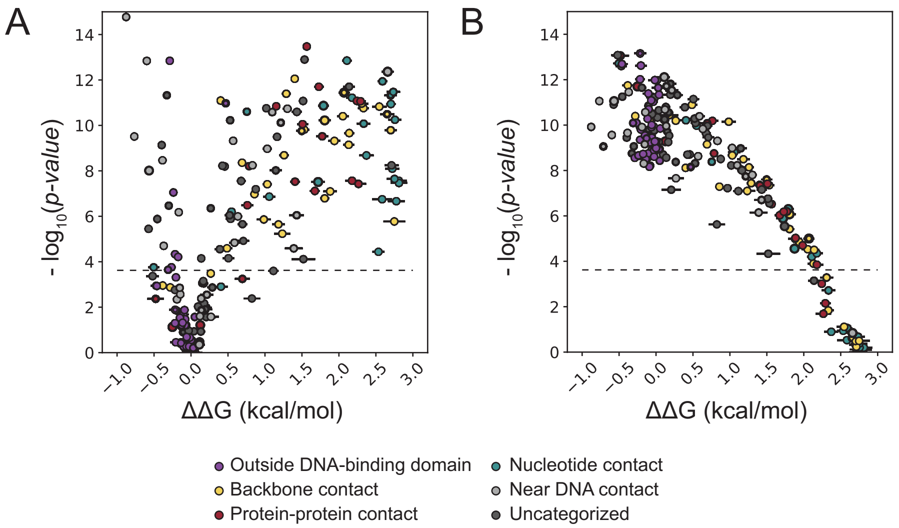


**Figure S6 (Related to Figure 3).** Determining dynamic range of assay measurements for reference sequence 5'- C CACGTG A -3'. **(A)** Comparison of the distribution of measured ΔΔGs (± SEM) for a given mutant *vs.* the distribution of ΔΔGs for WT Pho4 to determine statistically significant differences in binding (using a two-tailed t-test with a Bonferroni correction). Dashed line indicates estimated threshold for significance ($p$ = 0.00023). **(B)** Comparison of the distribution of measured ΔΔGs (± SEM) for a given mutant *vs.* the distribution of ΔΔGs for an inactive Pho4 mutant (A299D) to determine mutants resolvable from background noise. Dashed line indicates estimated threshold for significance ($p$ = 0.00023).

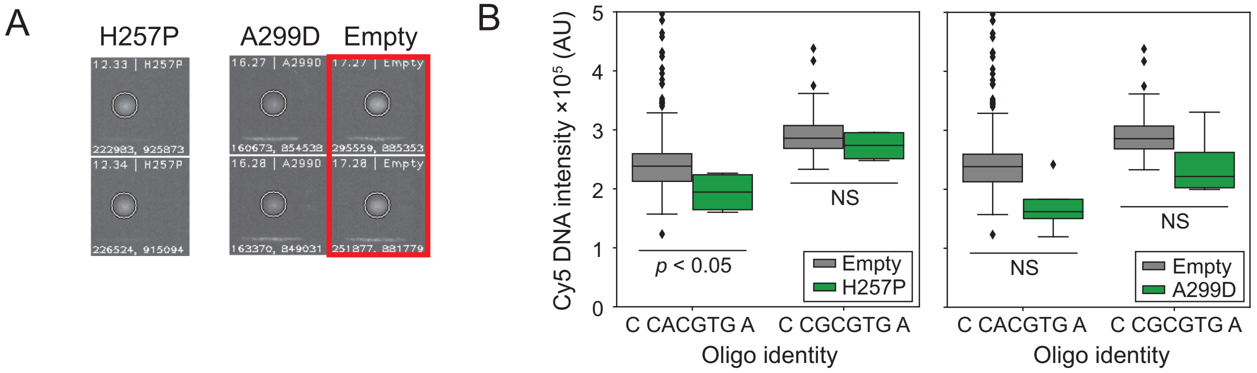


**Figure S7 (Related to Figure 3 and Methods).** Determining mutants at lower limit of assay detection. **(A)** Images showing measured DNA intensities for two Pho4 variants (H257P and A299D) and empty chambers (boxed in red) to determine effective lower limit of detection for binding to the 5'- C CACGTG A -3' reference sequence. Images are contrast-treated for visibility. **(B)** Distribution of raw Cy5 intensities at the highest DNA concentration for 2 binding deficient mutants (H257P and A299D) across 2 representative devices.

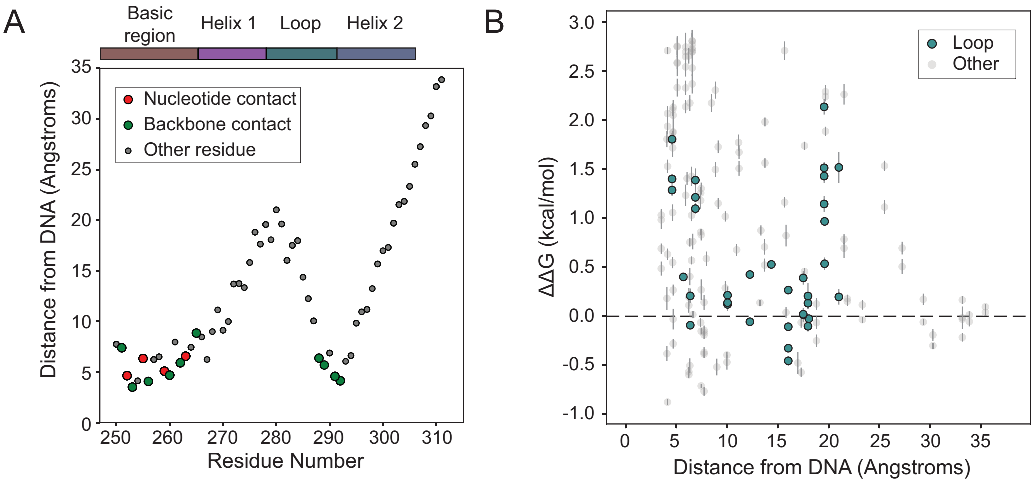


**Figure S8 (Related to Figure 3).** Measured effect of mutations on DNA affinity as a function of distance from DNA. **(A)** Distance (angstroms) of alpha carbon for each Pho4 residue from DNA, with DNA contacting residues annotated. **(B)** Measured relative change in binding energy (ΔΔG) as a function of distance from DNA for all Pho4 mutants; loop residues are shown in green.

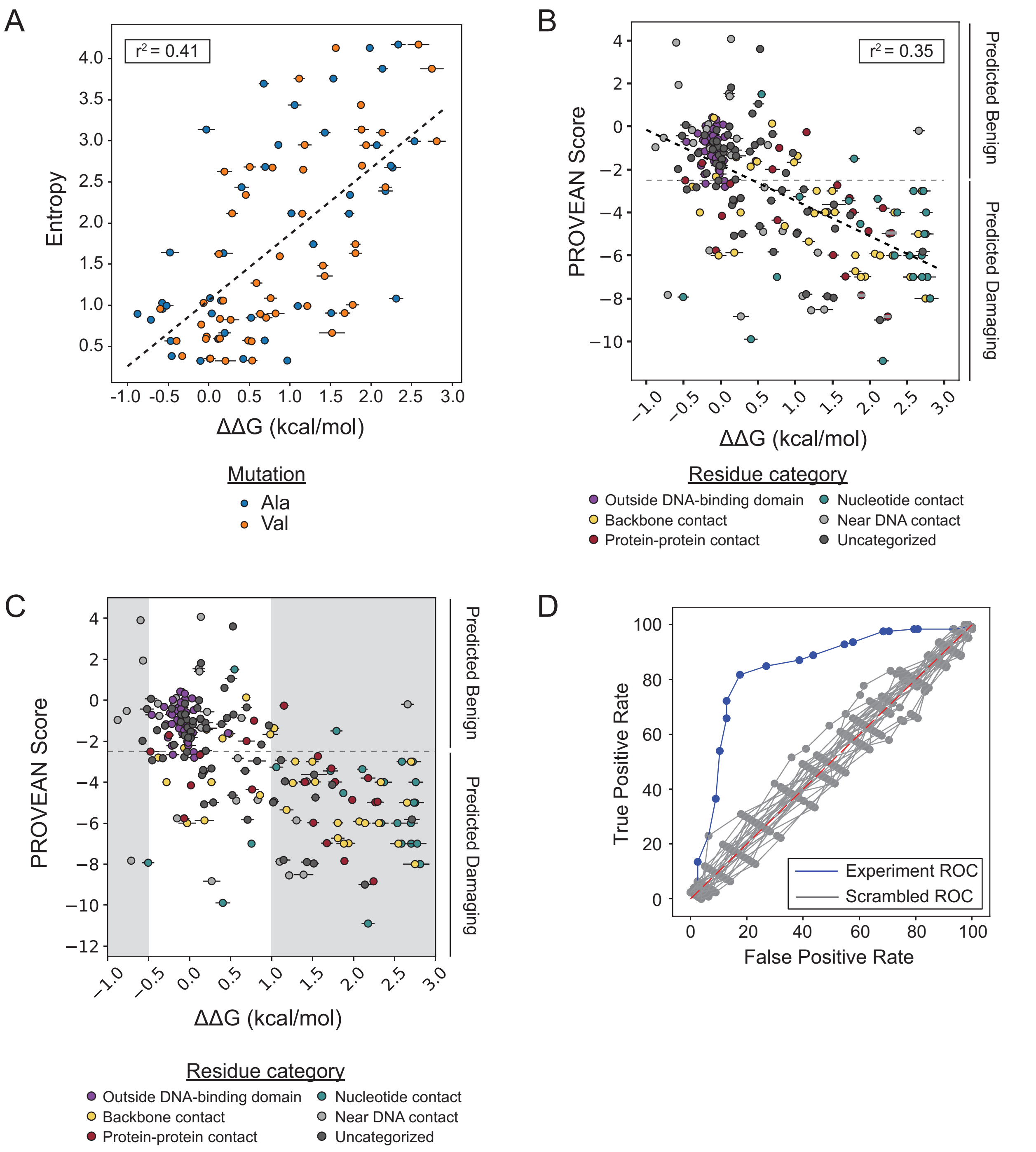


**Figure S9 (Related to Figure 3)**. Comparing observed effects of mutations with those predicted by phylogeny. **(A)** Entropy at a given residue position calculated based on PFAM alignment *vs.* measured mutational effects on affinity (median ΔΔG ± SEM). Dashed line indicates a linear fit between data points ($r^2$=0.41; y = 0.8*x* + 1.06). **(B)** Calculated PROVEAN score *vs*. measured mutational effect on affinity (median ΔΔG ± SEM). Horizontal dashed line indicates binding affinity of wildtype Pho4; diagonal dashed line indicates a linear regression of the data ($r^2$=0.35; y = -1.65*x* - 1.8). **(C)** Comparison of PROVEAN score *vs.* measured mutational effect on affinity (median ΔΔG ± SEM) for ROC analysis. Gray regions indicate threshold areas considered 'damaging' (ΔΔG < -0.5 kcal/mol or ΔΔG > 1 kcal/mol). Horizontal dashed line indicates standard damage threshold for PROVEAN. **(D)** ROC curve generated by adjusting damage threshold and calculating false positive and false negative rates. Blue curve indicates experimental ROC (AUC = 0.7). Gray curves indicate 25 separate trials of scrambled ΔΔG (AUC range = 0.43-0.55). Red dash line indicates AUC = 0.5 (random guess).

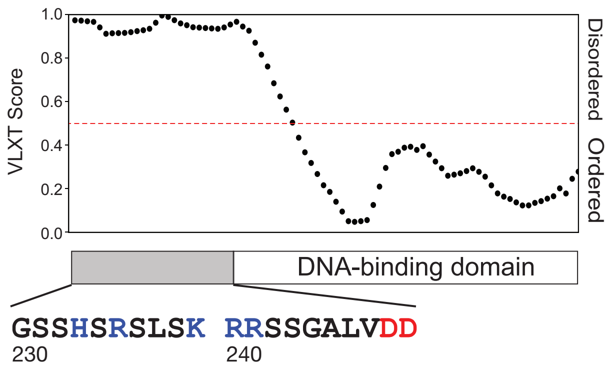


**Figure S10 (Related to Figure 3).** Predicted disorder of amino acids immediately preceding DNA binding domain of Pho4 (Obradovic, et al., 2003).

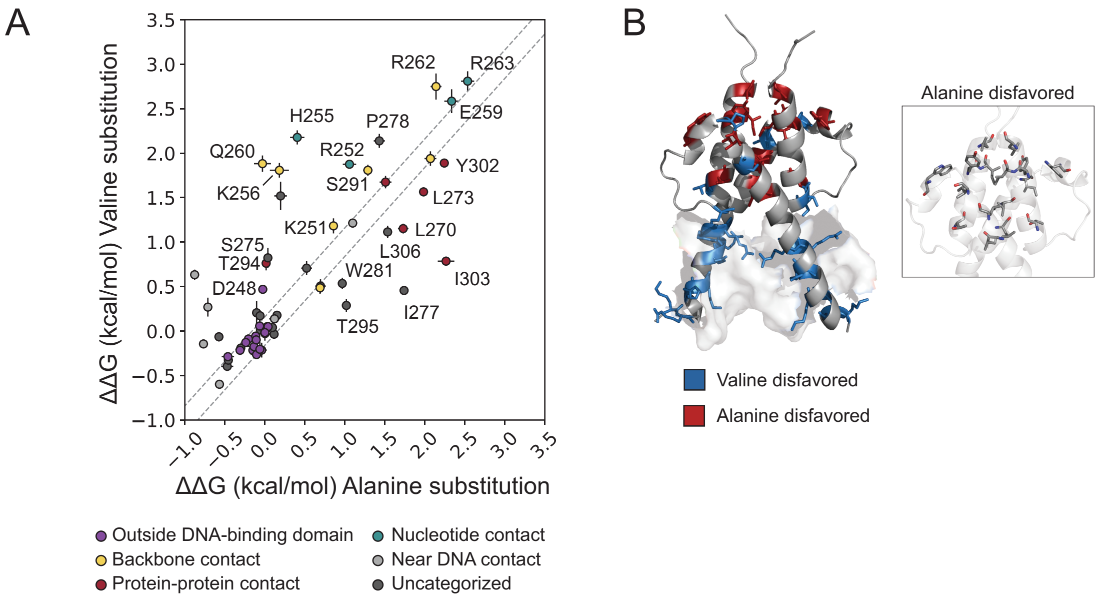


**Figure S11 (Related to Figure 3).** Differential effects of alanine and valine substitutions. **(A)** Measured ΔΔG values for valine *vs.* alanine substitutions at the same residue position; dashed lines indicate ±1 standard deviation of all wildtype Pho4 ΔΔG measurements. **(B)** Positions at which either alanine or valine are disfavored projected onto Pho4 crystal structure and alanine disfavored residues shown on transparent structure for clarity (inset).

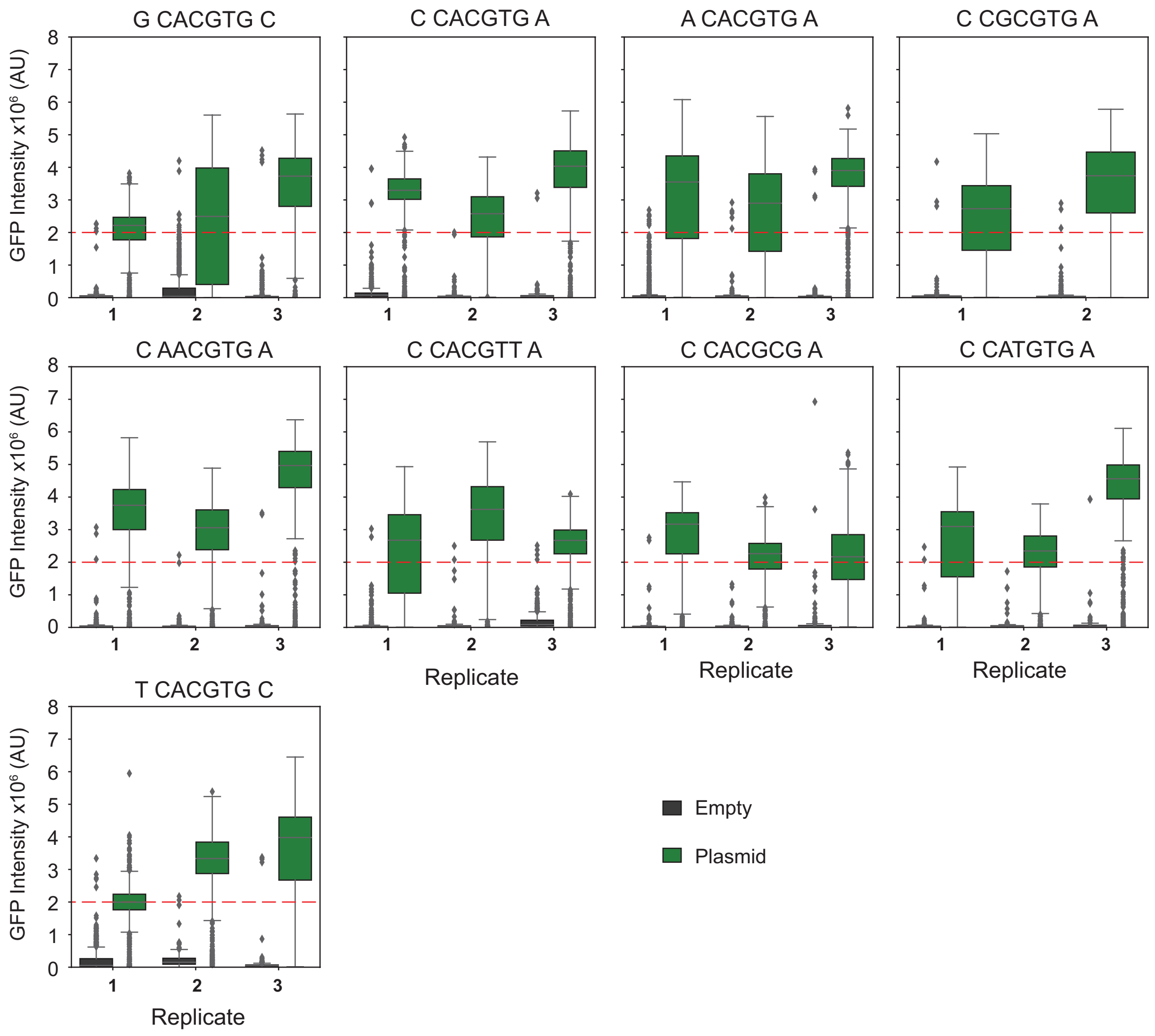


**Figure S12 (Related to Figure 4 and supplemental methods).** Measured TF GFP intensities for all experimental replicates for mutated DNA sequences in this study.

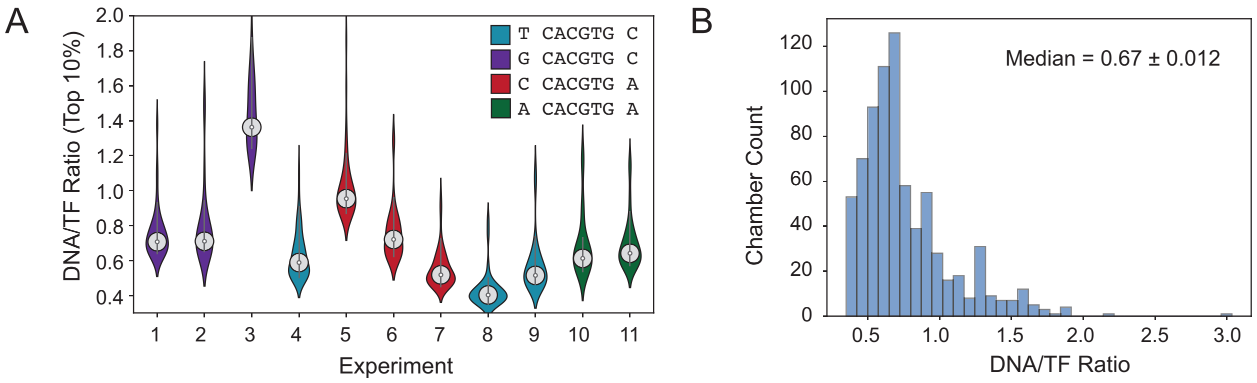


**Figure S13 (Related to Figure 4 and supplemental methods). (A)** Violin plots for the highest 10% of measured DNA/protein ratios at the highest measured DNA concentration for all experiments containing cognate E-box motif. **(B)** Distribution of number of chambers with a given DNA/TF ratio; median ratio = 0.67 ± 0.012 (SEM).

### G CACGTG C

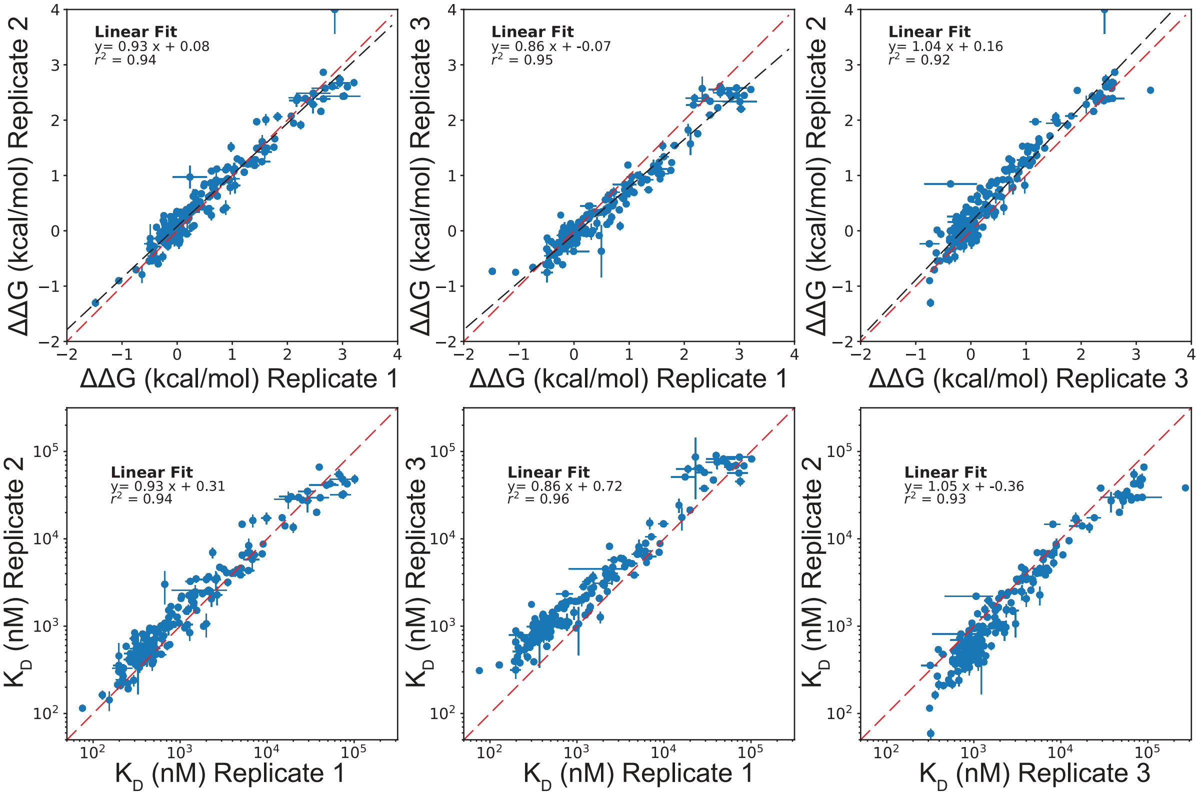


### T CACGTG C

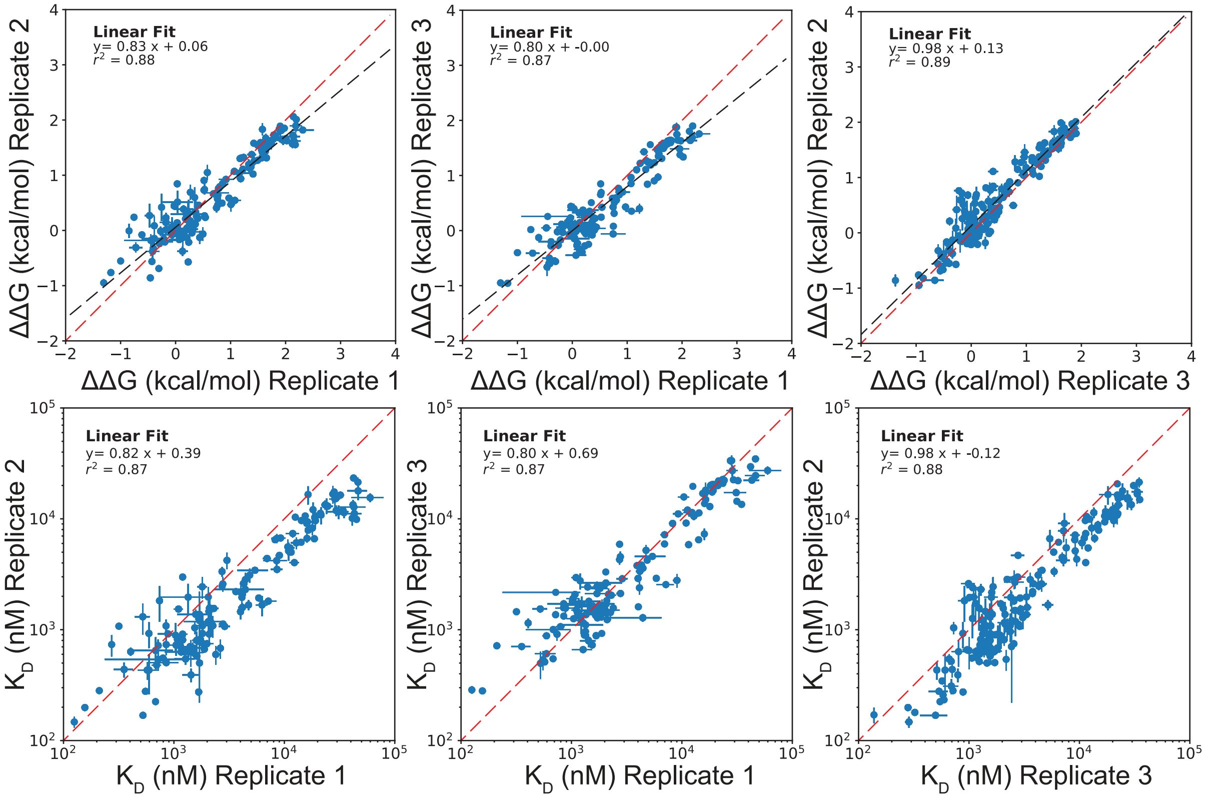

A CACGTG A

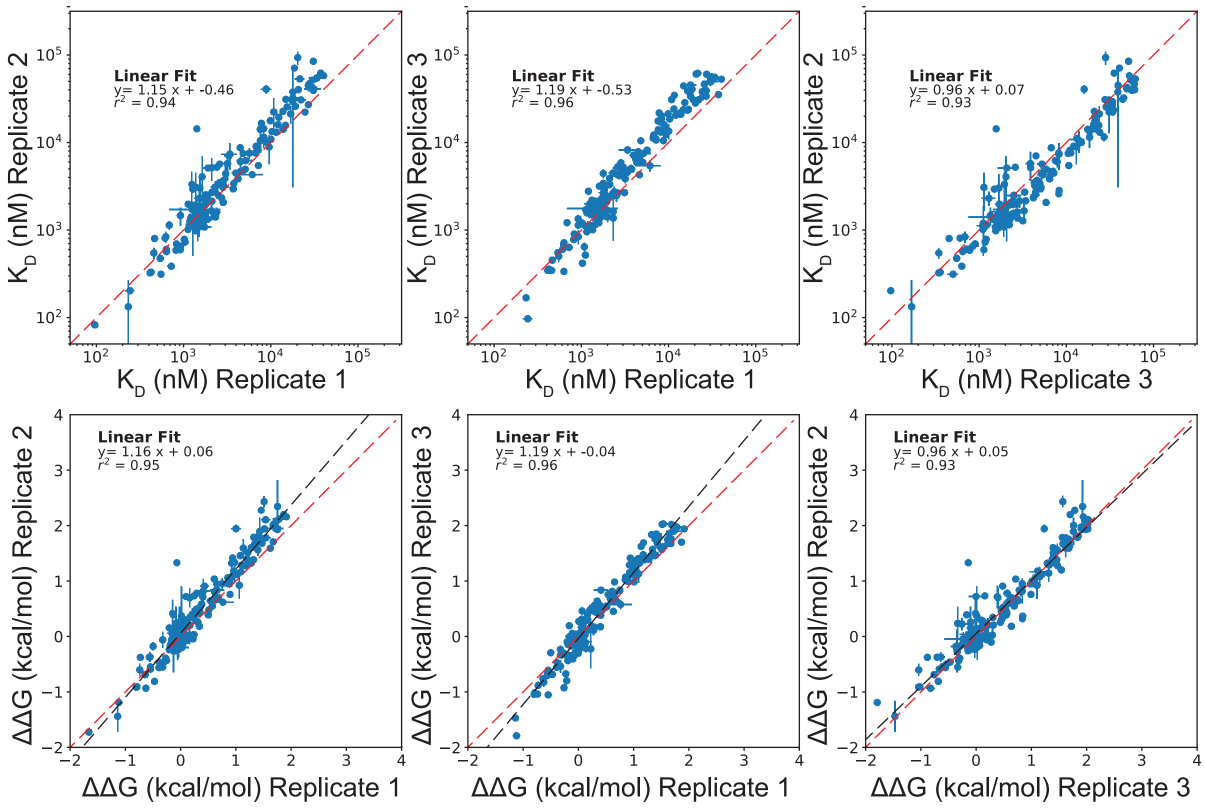


C AACGTG A

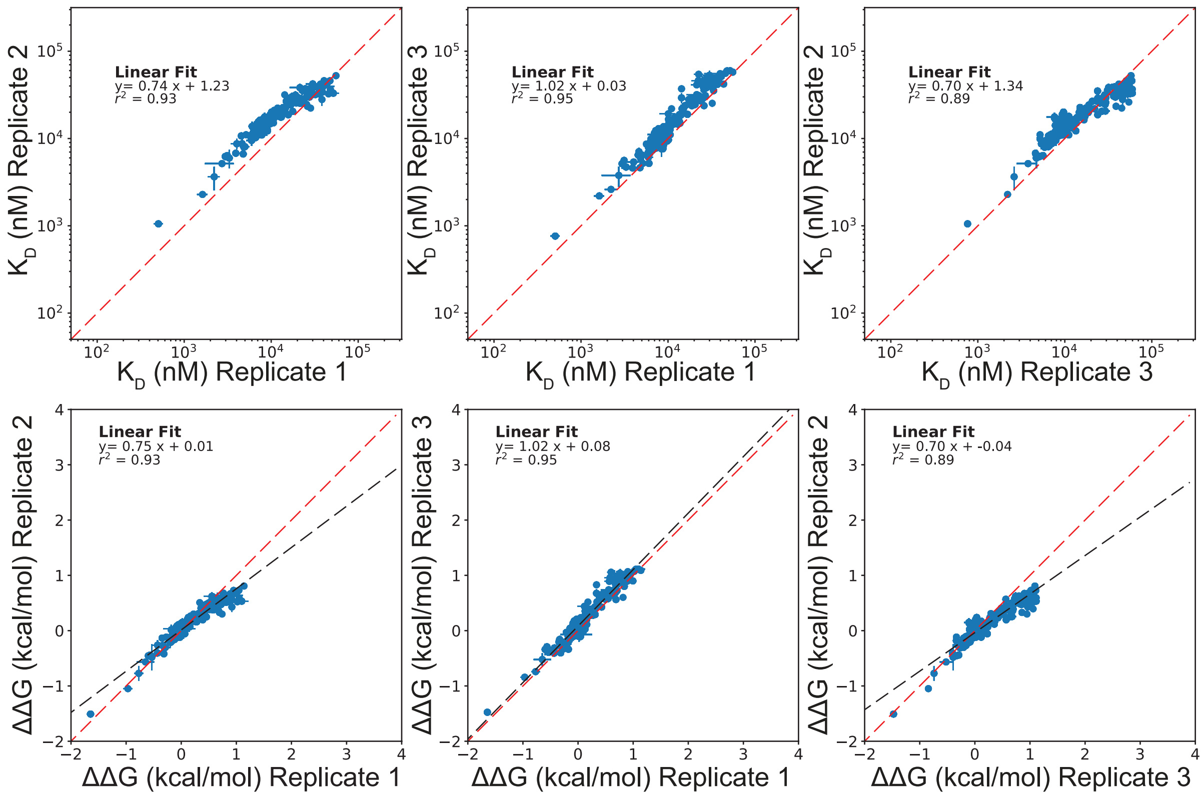

### C CACGTT A

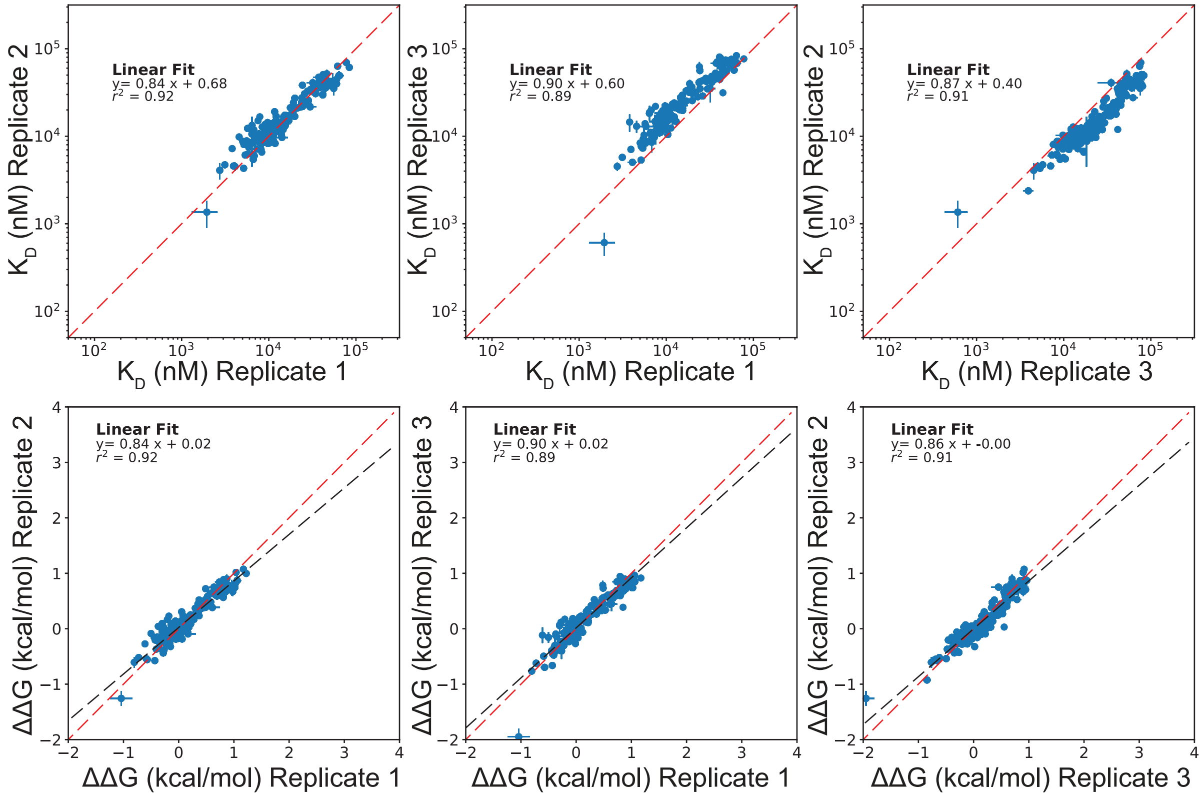


### C CACGCG A

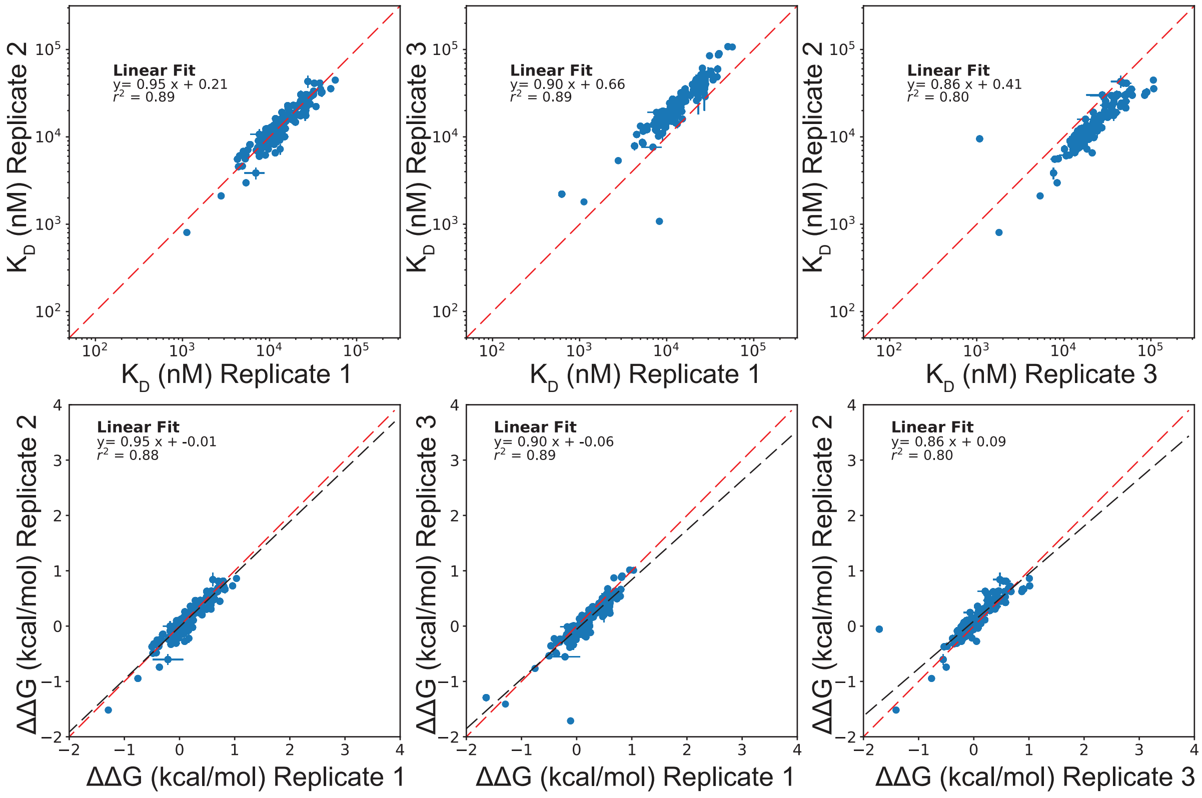

C CATGTG A

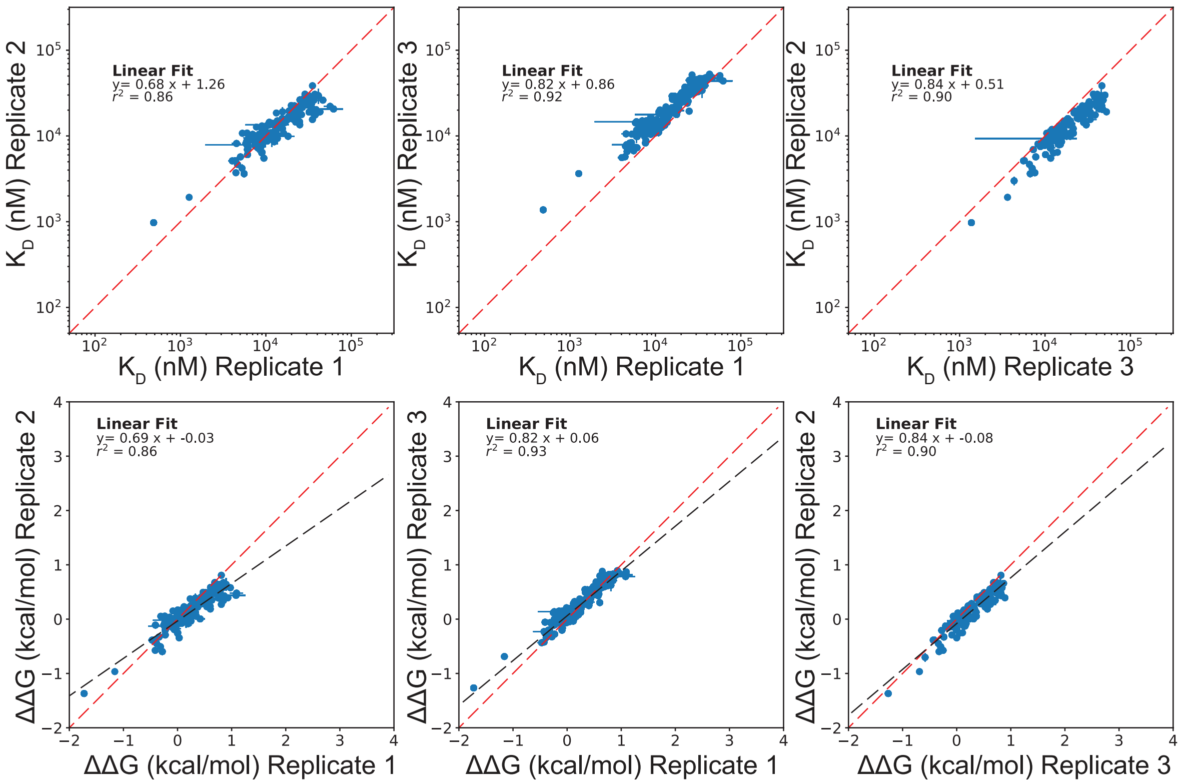


C CGCGTG A

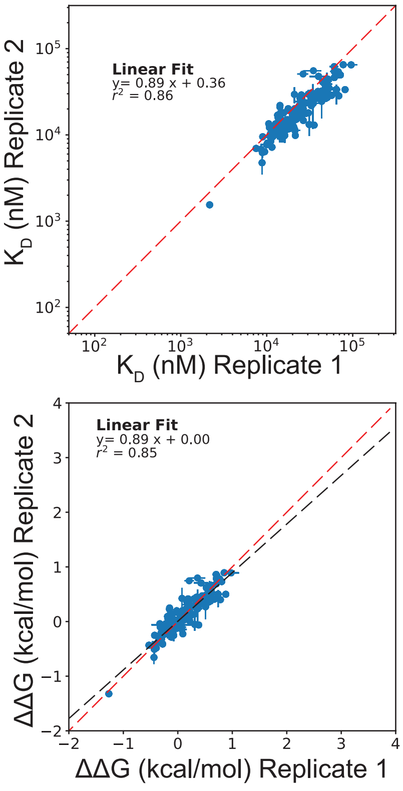

**Figure S14 (Related to Figure 4).** Pairwise comparison of per-mutant $K_d$s (top row) and ΔΔGs (bottom row) for all TF mutants for all replicates across all DNA sequences in this study. Points indicate median affinities (± SEM) for each TF mutant; all ΔΔGs are calculated relative to the WT Pho4 variant on a per-experiment basis. Black and red dashed lines indicate linear fits and identity lines, respectively.

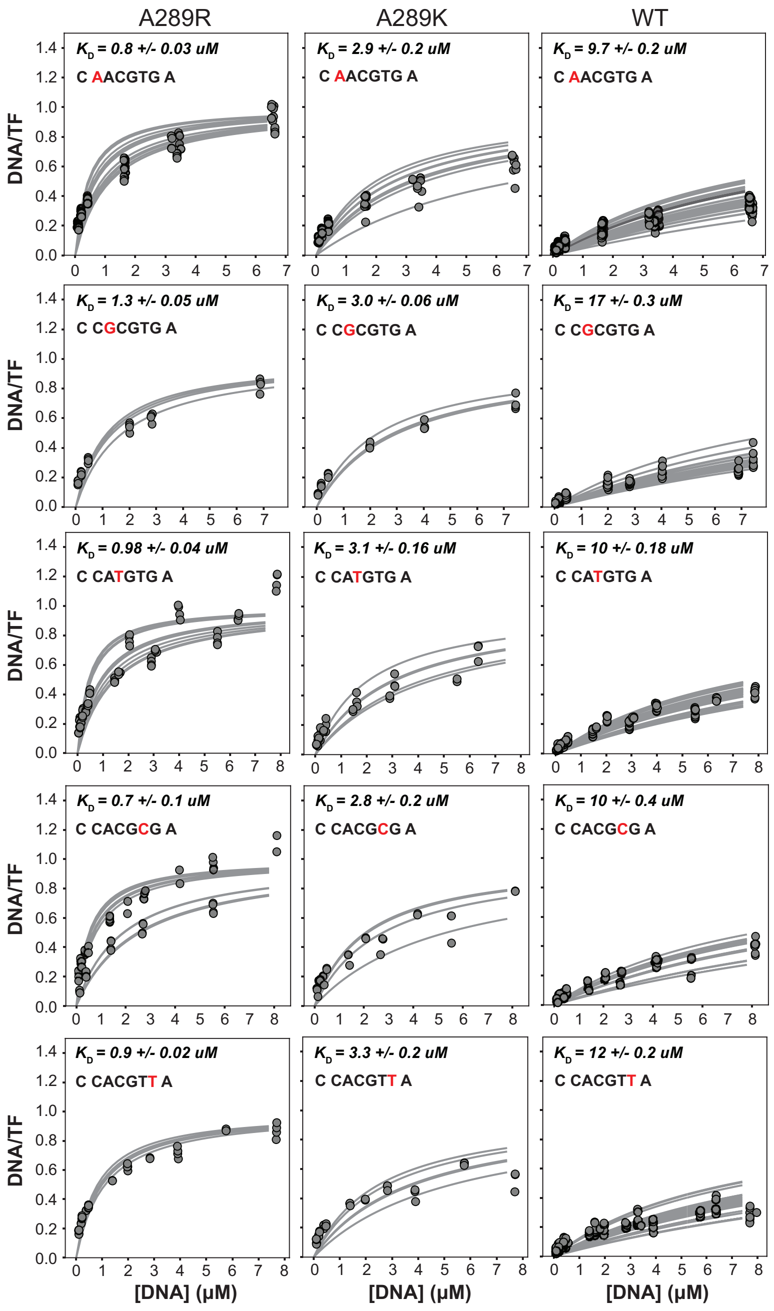


**Figure S15.** Concentration-dependent binding curves for mutations at the backbone-contacting residue A289 across all oligonucleotide sequences containing core mutations. A289R and A289K both increase binding affinities for all DNA sequences; $K_d$ values represent the median of all curves ± SEM.

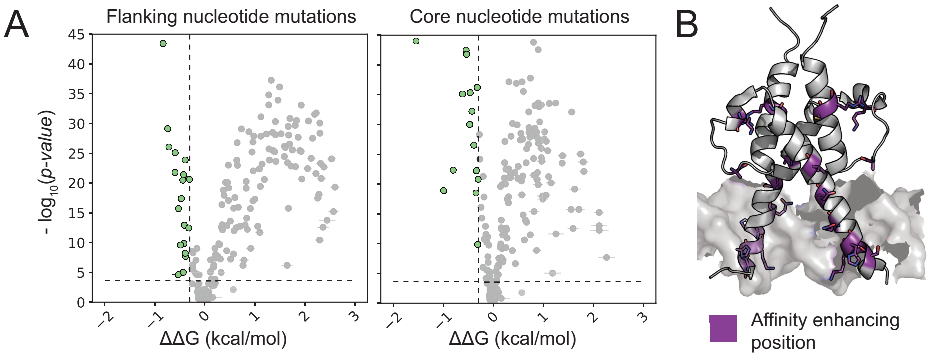


| Mutation category | Pho4 mutant identity |
|---|---|
| Outside DBD | H233A, H233V |
| Charge altering | D250Q, D250K, D250A, H255R, K256R, Q260R, K282A, K282V, A289K, A289R, R301A |
| Solvent-facing mutations | H257A, H257V, H271A, H271V, N264A, S254A, V268A |
| Other | C300A |

**Figure S16 (Related to Figure 4 and methods).** Affinity-enhancing Pho4 mutations. **(A)** Calculated -$\log_{10}$(*p*-values) (two-tailed T-test) *vs.* measured ΔΔG values (median ± SEM) for Pho4 variants interacting with DNA sequences containing mutations to flanking (left) or core (right) nucleotides; mutations with significantly enhanced binding are highlighted in green. **(B)** Location of affinity-enhancing mutations on Pho4 crystal structure and list of residues.

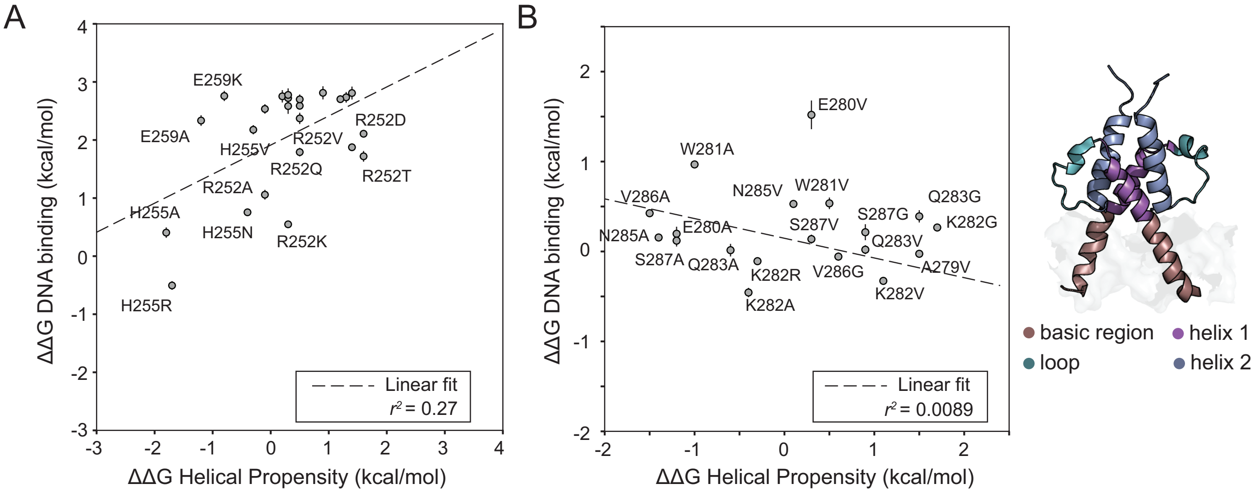


**Figure S17 (Related to Figure 5).** Measured ΔΔGs for substitutions to nucleotide-contacting and loop residues do not correlate with predicted changes in helical propensity. **(A)** Measured ΔΔGs for mutations at nucleotide-contacting residues *vs.* predicted changes in helical propensity ($r^2$ = 0.27; RMSE = 2.0 kcal/mol); as expected, ΔΔG effects are dominated by changes to residue/DNA contacts. **(B)** Measured ΔΔGs for mutations in unstructured loop region *vs.* predicted changes in helical propensity ($r^2$ = 0.0089; RMSE = 1.2 kcal/mol); as expected, ΔΔG effects in unstructured regions do not correlate with changes in helical propensity.

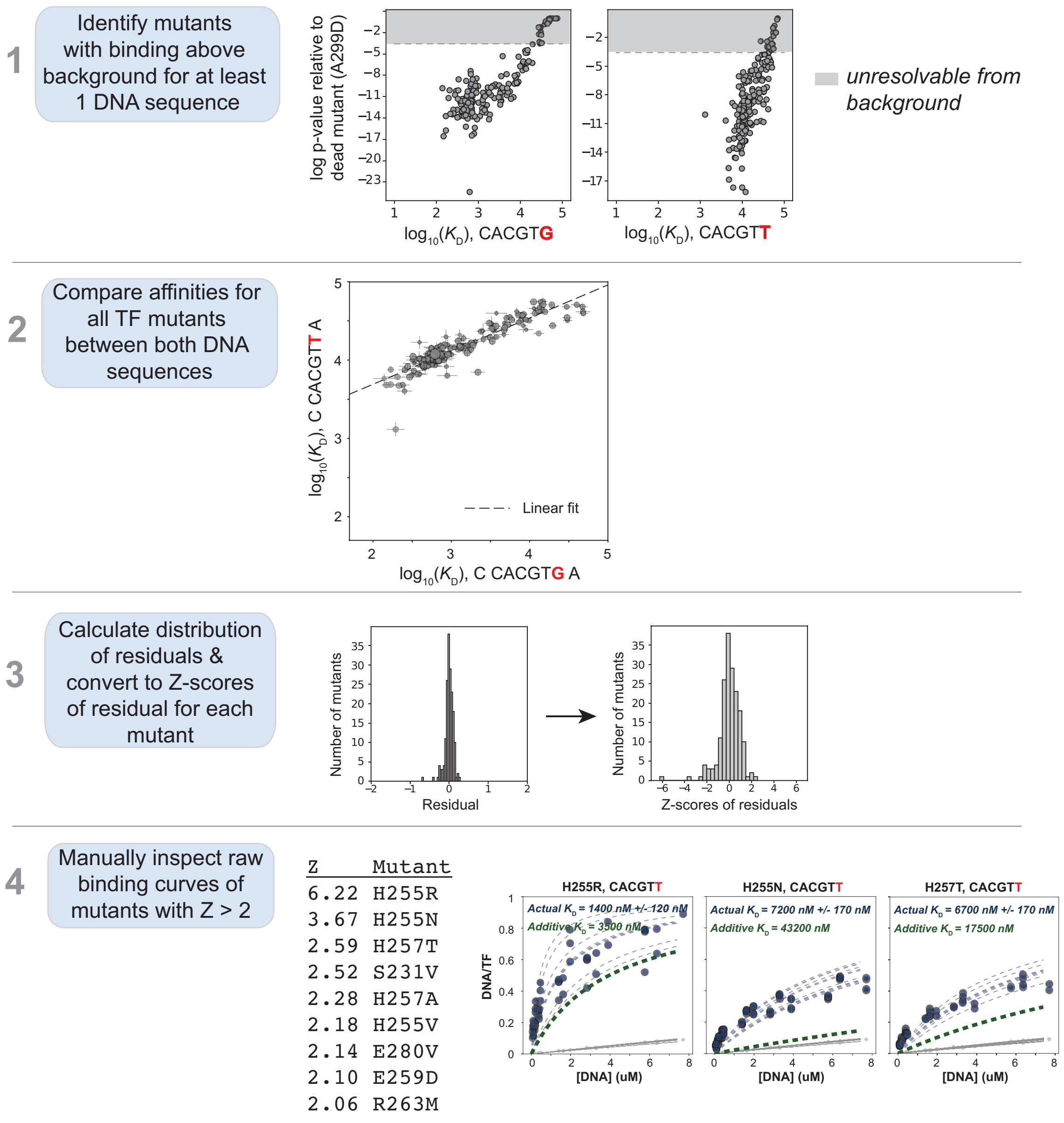


**Figure S18 (Related to Figure 6).** Analysis pipeline for identifying epistasis (non-additivity) between TF and DNA mutations.

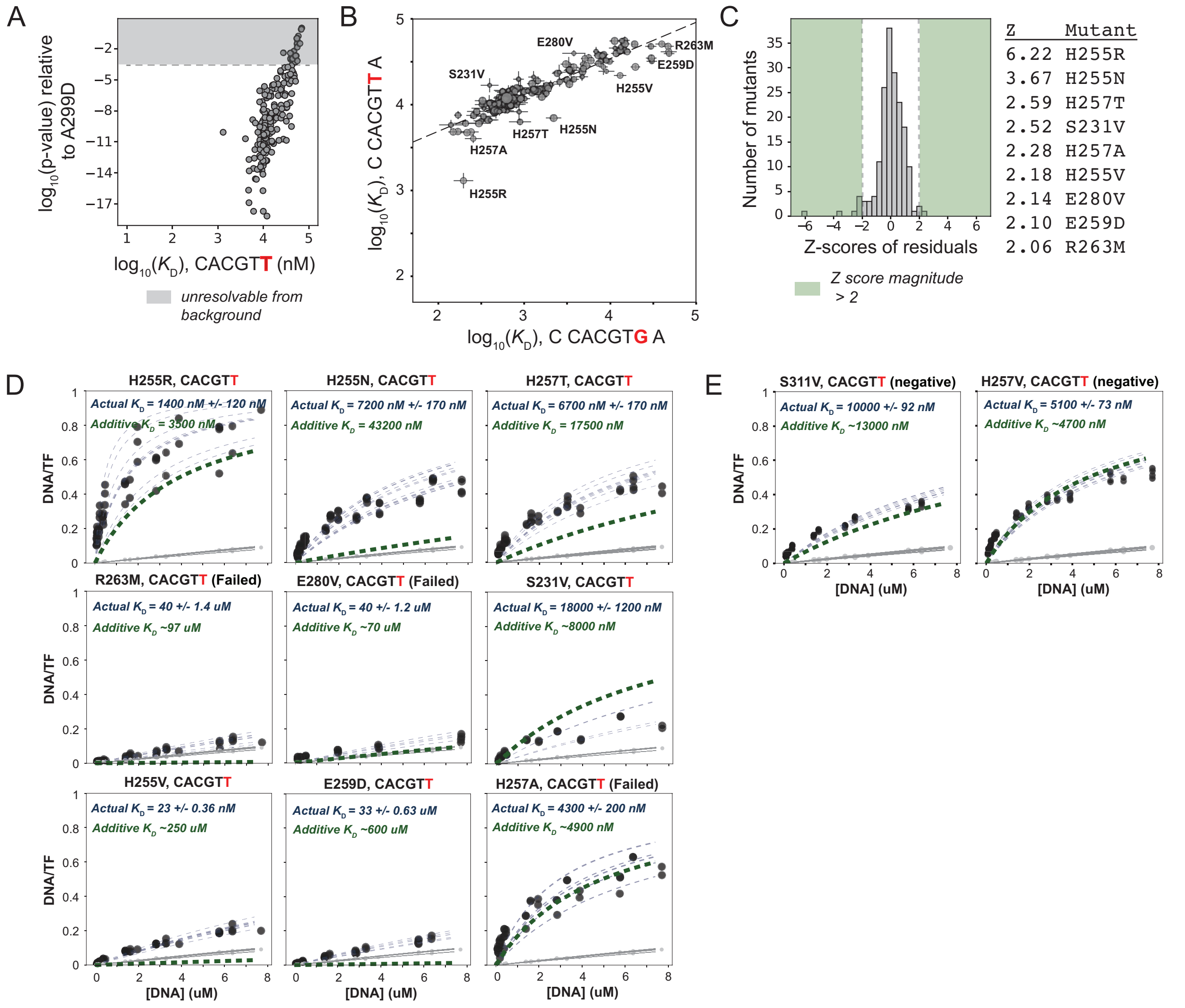


**Figure S19 (Related to Figure 6).** Double mutant cycle analysis across the TF-DNA interface for the 5'-C CACGTG A-3' and 5'-C CACGTT A-3' oligonucleotides. **(A)** Identification of mutants statistically resolvable from background for 5'-C CACGTT A-3' binding measurements. **(B)** Pairwise comparisons of measured $\log_{10}(K_d)$ values (median ± SEM) for Pho4 mutants for 5'-C CACGTG A-3' and 5'-C CACGTT A-3' oligonucleotides with outliers labeled. Dashed line indicates linear fit to plotted data ($r^2 = 0.84$). **(C)** Distribution of residual Z-scores (standardized residuals) with magnitude Z-score ≥ 2 highlighted and candidate mutants listed. **(D)** Validation of mutants with Z-scores ≥ 2 with failed mutants noted. **(E)** Example additive mutants conforming to non-epistatic expectation.

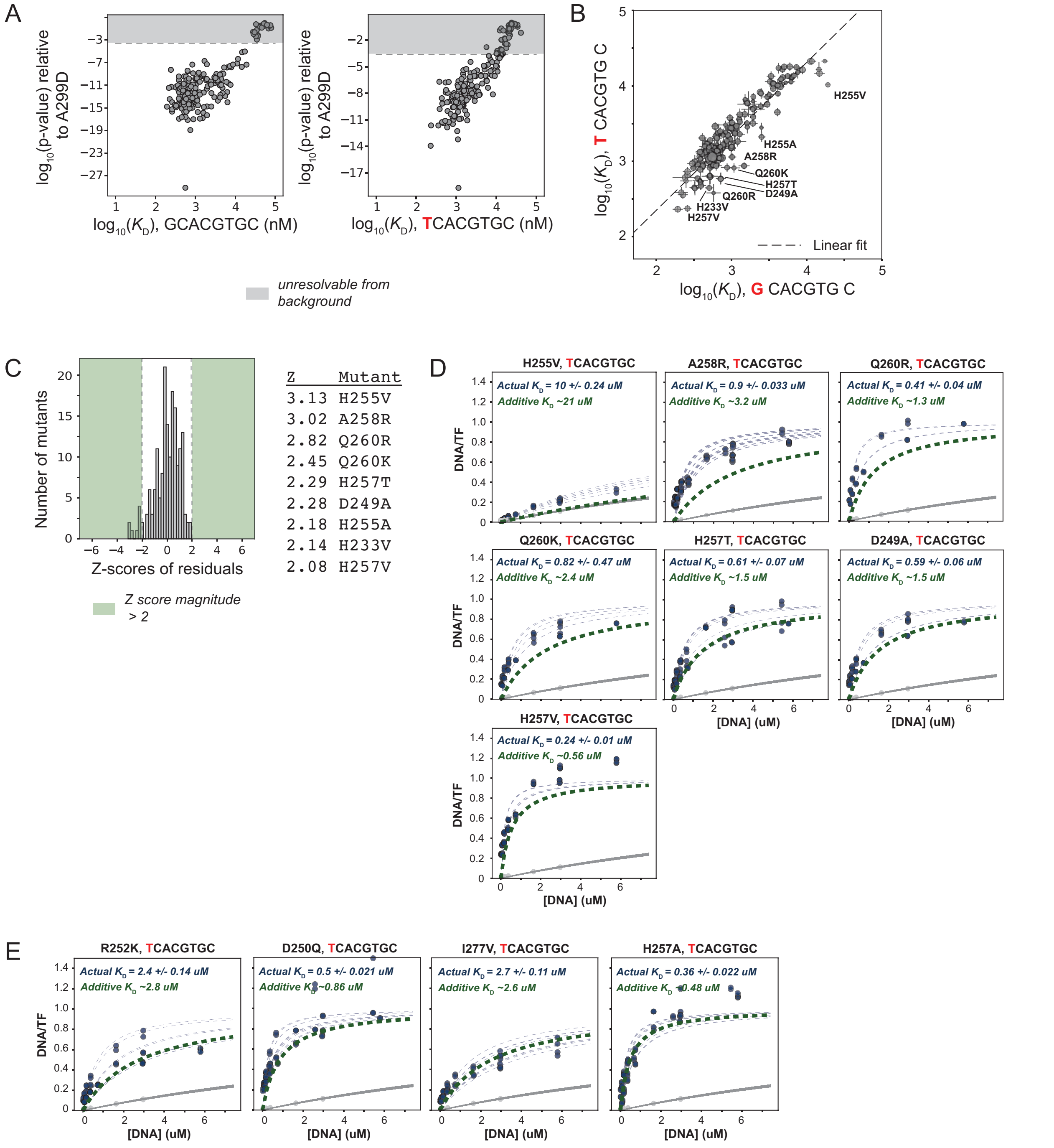


**Figure S20 (Related to Figure 6).** Double mutant cycle analysis across the TF-DNA interface for the 5'-G CACGTG C-3'and 5'-T CACGTG C-3 oligonucleotides. **(A)** Identification of mutants with binding statistically resolvable from background for 5'-G CACGTG C-3' and 5'-T CACGTG C-3' sequences. **(B)** Pairwise comparisons of measured $K_d$s (median ± SEM) for TF variants binding 5'-G CACGTG C-3' and 5'-T CACGTG C-3' DNA sequences with outliers marked; dashed line indicates linear fit ($r^2$ = 0.85). **(C)** Distribution of residual Z-scores (standardized residuals) with magnitude Z-score ≥ 2 highlighted and candidate mutants listed. **(D)** Validation of mutants with Z-scores ≥ 2 with failed mutants noted. **(E)** Example additive mutants conforming to non-epistatic expectation.

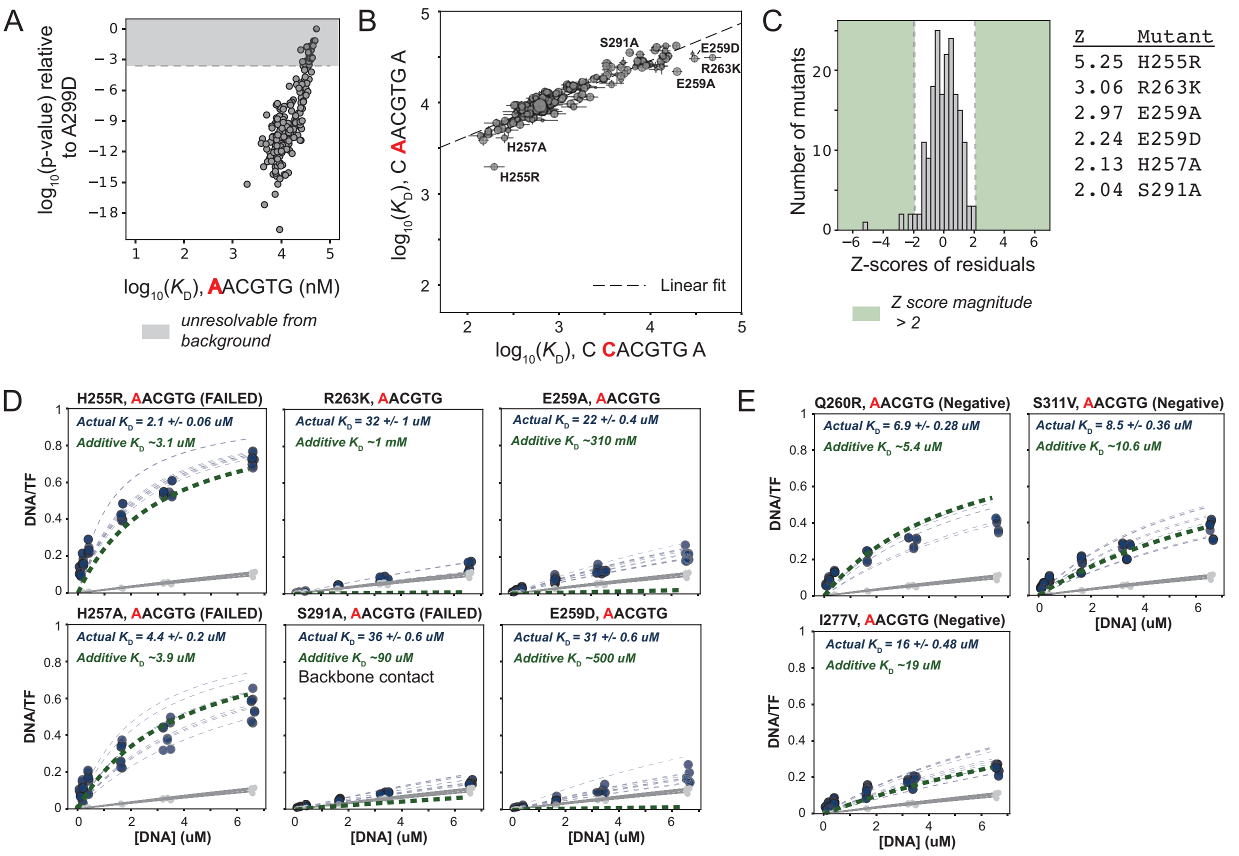


**Figure S21 (Related to Figure 6).** Double mutant cycle analysis across the TF-DNA interface for the 5'-G CACGTG C-3'and 5'-C AACGTG C-3 oligonucleotides. **(A)** Identification of mutants statistically resolvable from background for 5'-C AACGTG A-3' binding measurements. **(B)** Pairwise comparisons of measured affinities ($K_d$s, median ± SEM) for TF variants binding 5'-C CACGTG A-3' and 5'-C AACGTG A-3' DNA sequences with outliers marked; dashed line indicates linear fit ($r^2$ = 0.88). **(C)** Distribution of residual Z-scores (standardized residuals) with magnitude Z-score ≥ 2 highlighted and candidate mutants listed. **(D)** Validation of mutants with Z-scores ≥ 2 with failed mutants noted. **(E)** Example additive mutants conforming to non-epistatic expectation.

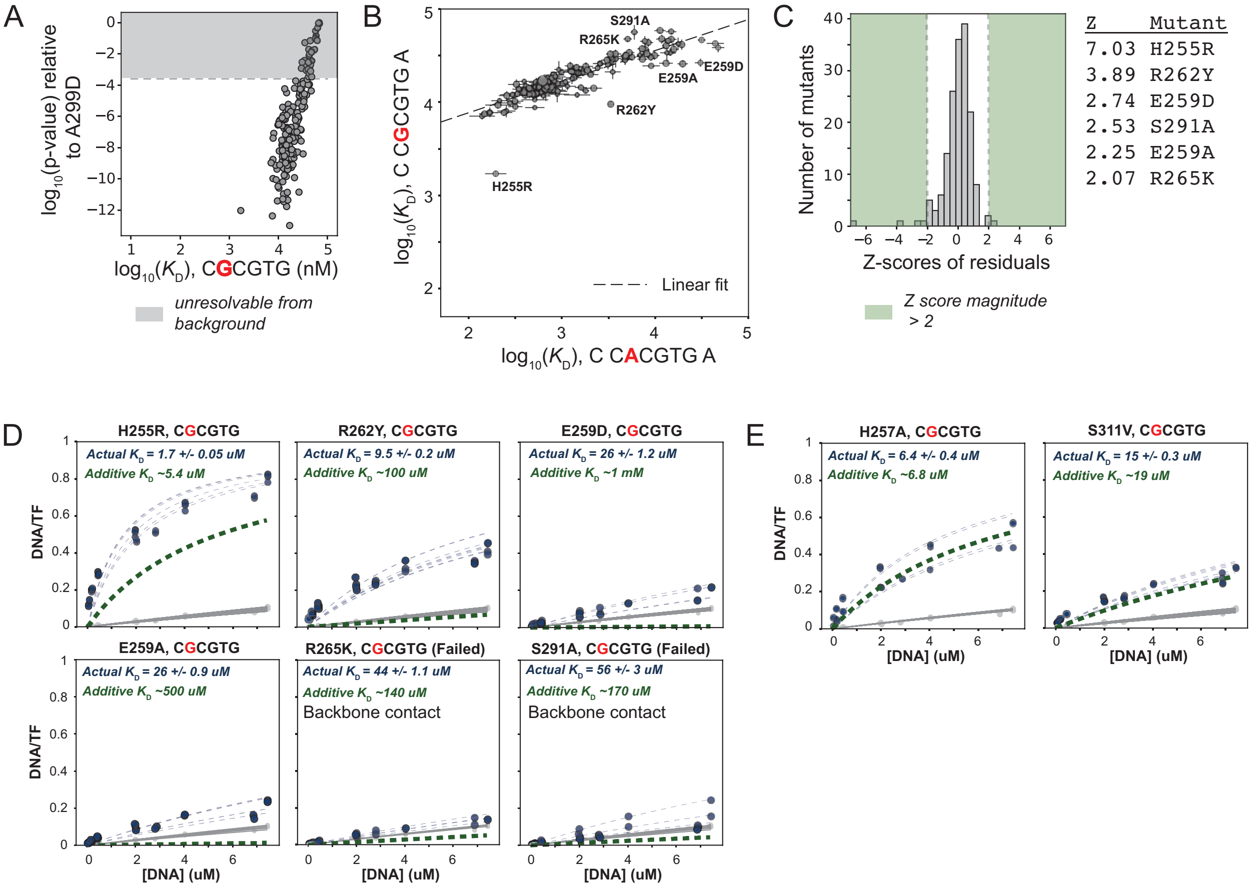


**Figure S22 (Related to Figure 6).** Double mutant cycle analysis across the TF-DNA interface for the 5'-G CACGTG C-3'and 5'-C CGCGTG C-3 oligonucleotides. **(A)** Identification of mutants statistically resolvable from background for 5'-C CGCGTG A-3' binding measurements. **(B)** Pairwise comparisons of measured affinities ($K_d$s, median ± SEM) for TF variants binding 5'-C CACGTG A-3' and 5'-C CGCGTG A-3' DNA sequences with outliers marked; dashed line indicates linear fit ($r^2$ = 0.76). **(C)** Distribution of residual Z-scores (standardized residuals) with magnitude Z-score ≥ 2 highlighted and candidate mutants listed. **(D)** Validation of mutants with Z-scores ≥ 2 with failed mutants noted. **(E)** Example additive mutants conforming to non-epistatic expectation.

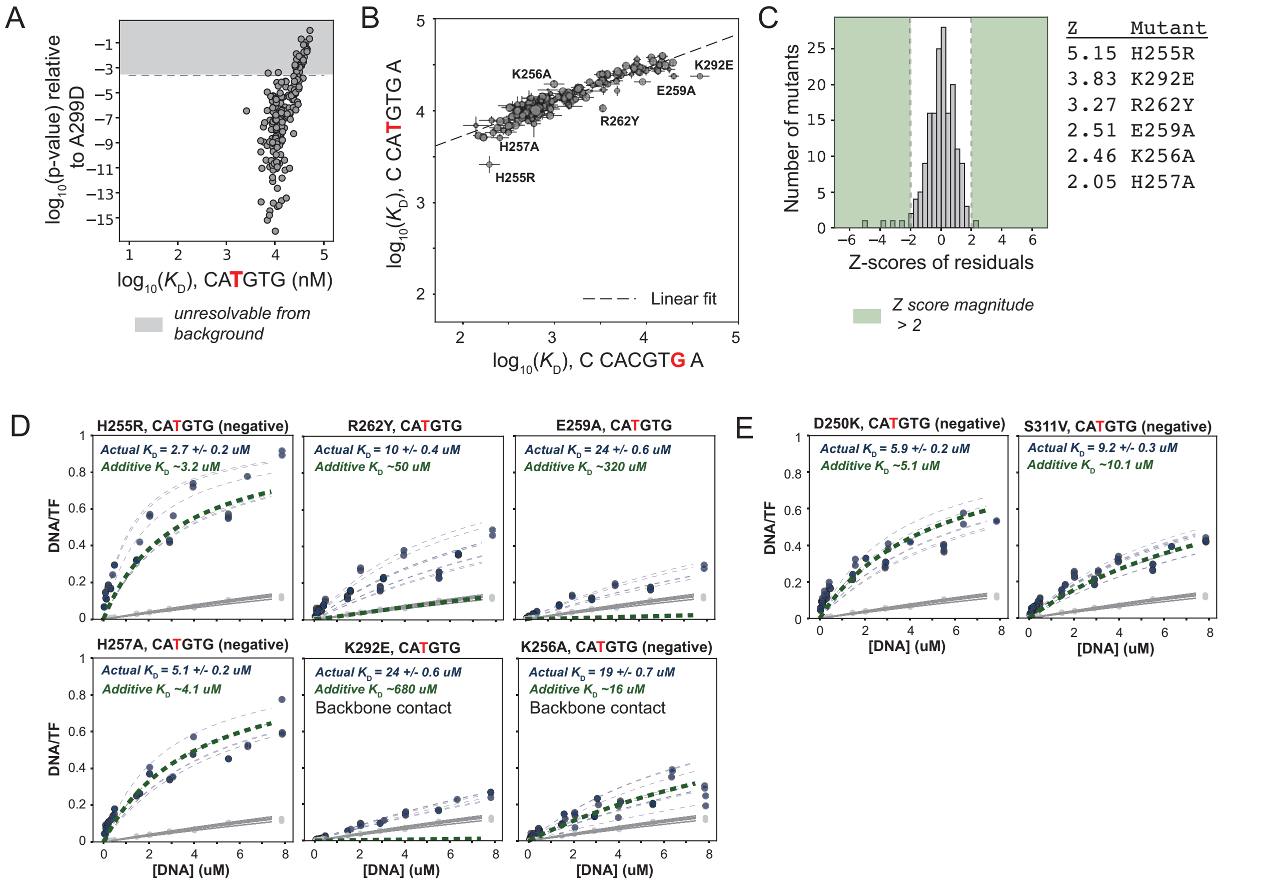


**Figure S23 (Related to Figure 6).** Double mutant cycle analysis across the TF-DNA interface for the 5'-G CACGTG C-3'and 5'-C CATGTG C-3 oligonucleotides. **(A)** Identification of mutants statistically resolvable from background for 5'-C CATGTG A-3' binding measurements. **(B)** Pairwise comparisons of measured affinities ($K_d$s, median ± SEM) for TF variants binding 5'-C CACGTG A-3' and 5'-C CATGTG A-3' DNA sequences with outliers marked; dashed line indicates linear fit to plotted data ($r^2 = 0.86$). **(C)** Distribution of residual Z-scores (standardized residuals) with magnitude Z-score ≥ 2 highlighted and candidate mutants listed. **(D)** Validation of mutants with Z-scores ≥ 2 with failed mutants noted. **(E)** Example additive mutants conforming to non-epistatic expectation.

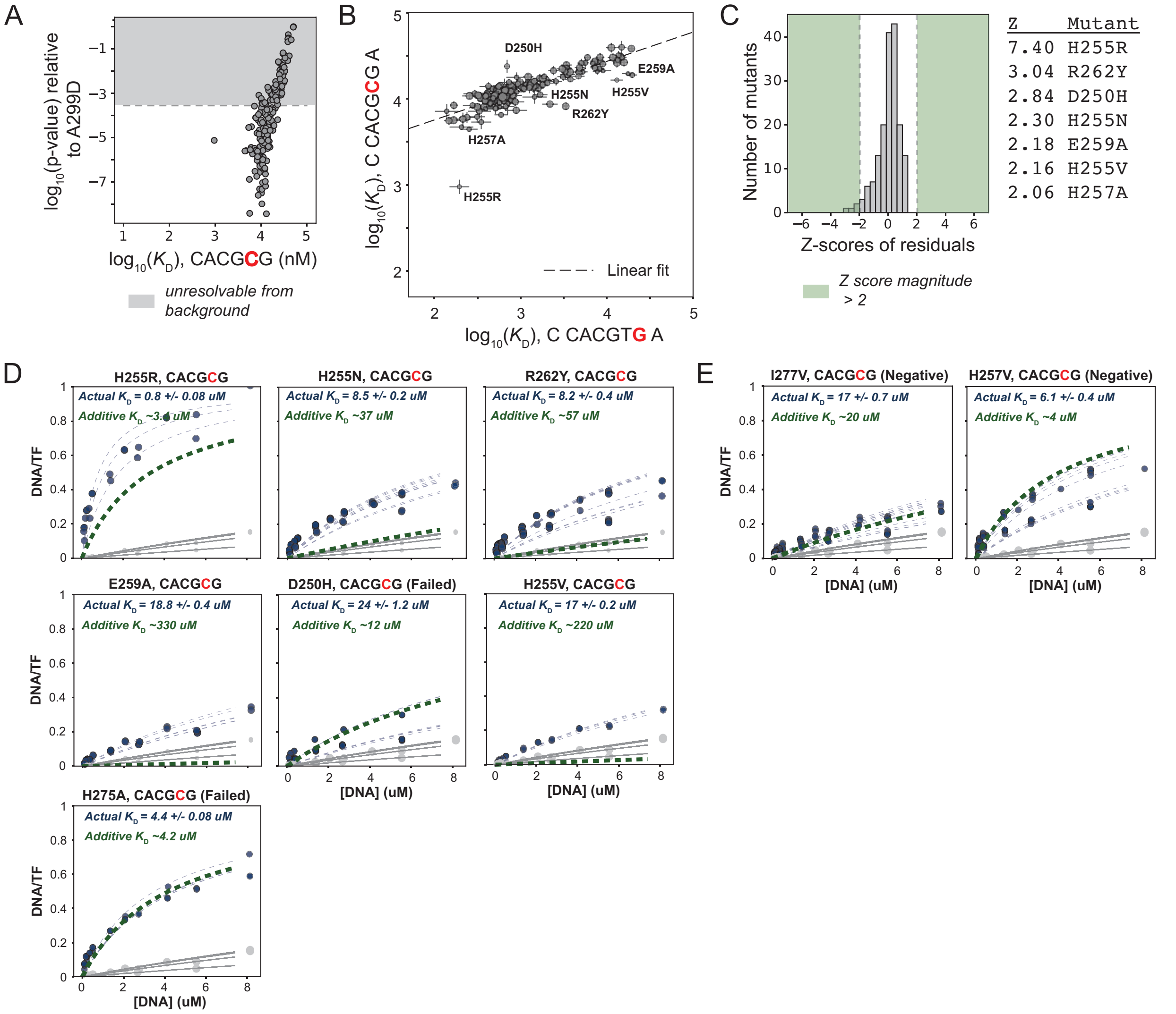


**Figure S24 (Related to Figure 6).** Double mutant cycle analysis across the TF-DNA interface for the 5'-G CACGTG C-3'and 5'-C CACGCG C-3 oligonucleotides. **(A)** Identification of mutants statistically resolvable from background for 5'-C CACGCG A-3' binding measurements. **(B)** Pairwise comparisons of median affinities (± SEM) for TF mutants between 5'-CCACGTGA-3' and 5'- CCACGCGA -3' DNA sequences with outliers marked. Dashed line indicates linear fit to plotted data. Linear fit $r^2 = 0.69$. ). **(C)** Distribution of residual Z-scores (standardized residuals) with magnitude Z-score ≥ 2 highlighted and candidate mutants listed. **(D)** Validation of mutants with Z-scores ≥ 2 with failed mutants noted. **(E)** Example additive mutants conforming to non-epistatic expectation.

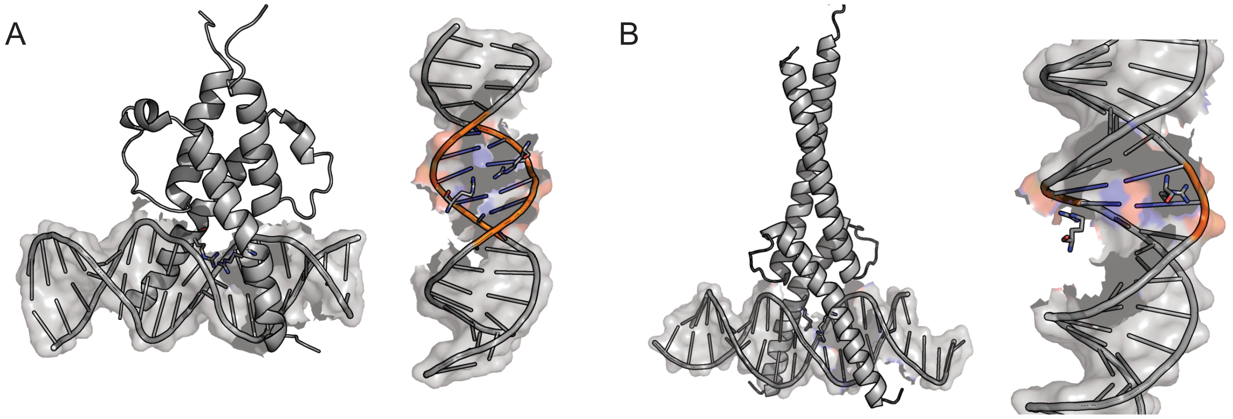

**Figure S25 (Related to Figure 6).** Comparison of 13$^{th}$ arginine in DNA binding domains for Pho4 and SREBP1. **(A)** Pho4 bound to DNA (PDB: 1A0A) (left); DNA molecule with 13$^{th}$ arginine residue in Pho4 (R263) shown as sticks making contacts across central nucleotides in E-box motif (right). Portion of DNA sequence containing E-box indicated by orange DNA backbone. **(B)** SREBP1 bound to DNA (PDB: 1AM9) (left); DNA molecule with 13$^{th}$ arginine residue in SREBP1 shown as sticks contacting DNA backbone (right). Conformations of arginine for both monomers are symmetric regardless of half-site identity. Portion of DNA sequence containing E-box indicated by orange DNA backbone.
